# Extrachromosomal Amplification of Human Papillomavirus Episomes as a Mechanism of Cervical Carcinogenesis

**DOI:** 10.1101/2021.10.22.465367

**Authors:** Nicole M. Rossi, Jieqiong Dai, Yi Xie, Hong Lou, Joseph F. Boland, Meredith Yeager, Roberto Orozco, Enrique Alvirez, Lisa Mirabello, Eduardo Gharzouzi, Michael Dean

**Author notes:** correspondence to: Dr. Michael Dean, 9615 Medical Center Drive, Rm 3130 Rockville, MD 20850. These authors contributed equally (Nicole M. Rossi, Jieqiong Dai, Yi Xie).

## Abstract

Integration of Human Papillomaviruses (HPV) is an important mechanism of carcinogenesis but is absent in a significant fraction of HPV16+ tumors. We applied long-read whole-genome sequencing (WGS) to cervical cancer cell lines and tumors. In two HPV16+ cell lines, we identified large tandem arrays of full-length and truncated viral genomes integrated into multiple locations indicating formation as extrachromosomal DNA (HPV superspreading). An HPV16+ cell line with episomal DNA has tandem arrays of full-length, truncated, and rearranged HPV16 genomes (multimer episomes). WGS of HPV16+ cervical tumors revealed that 11/20 with only episomal HPV (EP) have intact monomer episomes. The remaining nine EP tumors have multimer and rearranged HPV genomes. Most HPV rearrangements disrupt the *E1* and *E2* genes, and EP tumors overexpress the *E6* and *E7* viral oncogenes. Tumors with both episomal and integrated HPV16 display multimer episomes and concatemers of human and viral sequences. One tumor has a recurrent deletion of an inhibitory site regulating *E6* and *E7* expression, and another has a recurrent duplication consistent with HPV superspreading. Therefore, HPV16 can cause cancer without integration through aberrant episomal replication, forming rearranged and multimer episomes.

## INTRODUCTION

Human Papillomavirus (HPV) causes over 90% of cervical cancer, resulting in at least 300,000 deaths per year, mostly in low-and-middle-income countries (LMICs)(Schiffman et al., 2016; Walboomers et al., 1999). Two high-risk types, HPV16 and HPV18, are responsible for over 70% of precancerous cervical lesions and advanced cancers (Schiffman and Castle, 2005). However, genetic variation within HPV types is associated with the histological type of cervical cancer (adenocarcinoma versus squamous cell carcinoma (SCC)), integration rate, and carcinogenicity (Bodelon et al., 2016; Cancer Genome Atlas Research et al., 2017; Mirabello et al., 2016).

The HPV genome replicates inside the host cell as an episome, a form of extracellular DNA (ecDNA). The circular, 7900 base pair (bp) genome encodes two critical oncoproteins, E6 and E7, which inhibit the tumor suppressors TP53 and RB1 (Foster and Galloway, 1996; Scheffner et al., 1990; Werness et al., 1990). E6 and E7 oncogene expression is controlled in part by the E2 protein (Bellanger et al., 2011), and E1 and E2 regulate replication of the viral genome (Dowhanick et al., 1995). In most cancers, the viral genome integrates into the host DNA deleting all or portions of the *E1* and *E2* genes (Schwarz et al., 1985). Thus, most integration events retain a truncated part of the HPV genome containing the viral regulatory region and the *E6* and *E7* genes (type 1 integration), In contrast, a subset of tumors integrates intact copies of the viral genome (type 2 integration) (Jeon et al., 1995). Inactivation of E1 and E2 inhibits DNA synthesis at the HPV origin of replication and releases repression of E6 and E7 expression. HPV integration can be associated with amplified viral and flanking cellular genes at the integration site (Akagi et al., 2014; Baker et al., 1987; Parfenov et al., 2014; Peter et al., 2010; Wagatsuma et al., 1990) and lead to activation of flanking human genes (Warburton et al., 2018). Integration can occur throughout the genome but mainly occurs in transcriptionally active regions. Recurrent sites have been found in or near specific genes such as *MYC, TP63, FHIT, LRP1B, RAD51B, CDK13*, and *KLF5*/*KLF12 (Bodelon et al., 2016; Couturier et al., 1991; Hu et al., 2015; Parfenov et al., 2014; Peter et al., 2010; Tang et al., 2020)*.

Viral integration is not a part of the normal lifecycle of HPV and does not occur in all tumors. While HPV18 and HPV45 integrate into nearly 100% of tumors, the integration rate of HPV16 is only 60-80% (Bodelon et al., 2016; Dutta et al., 2015; Hafner et al., 2008; Hu et al., 2015; Lou et al., 2020; Vinokurova et al., 2008). Some episomal-only HPV infected tumors have mutations in the upstream regulatory region (URR) of HPV (Lace et al., 2009; May et al., 1994), and HPV16 regulation is altered in cellular models of episomal infection (Gray et al., 2010). To investigate the mechanism of HPV16 carcinogenesis in the absence of integration, we applied long-read DNA and RNA sequencing to cell lines with and without HPV integration. Finally, we used these methods to study a well-curated set of HPV16 tumors with episomal DNA. Our results provide insight into the extrachromosomal replication of HPV and carcinogenesis.

## Results

### HPV type controls integration frequency

Different HPV types have varying frequencies of integration (Bodelon et al., 2016; Wentzensen et al., 2004). To explore HPV integration frequency by HPV type, we compiled data from The Cancer Genome Atlas (TCGA) and Guatemalan tumors (Cancer Genome Atlas Research et al., 2017; Lou et al., 2020; Lou et al., 2015). We confirmed that HPV18 and HPV45 integrate into nearly 100% of tumors, while HPV16 and HPV31 only integrate into 60-70% of tumors (**Figure 1A**). Furthermore, HPV18 and 45 are in the alpha-7 clade and HPV16 and 31 the alpha-9 clade (Burk et al., 2013), indicating that viral genetics determines integration rate.

**Figure 1.**
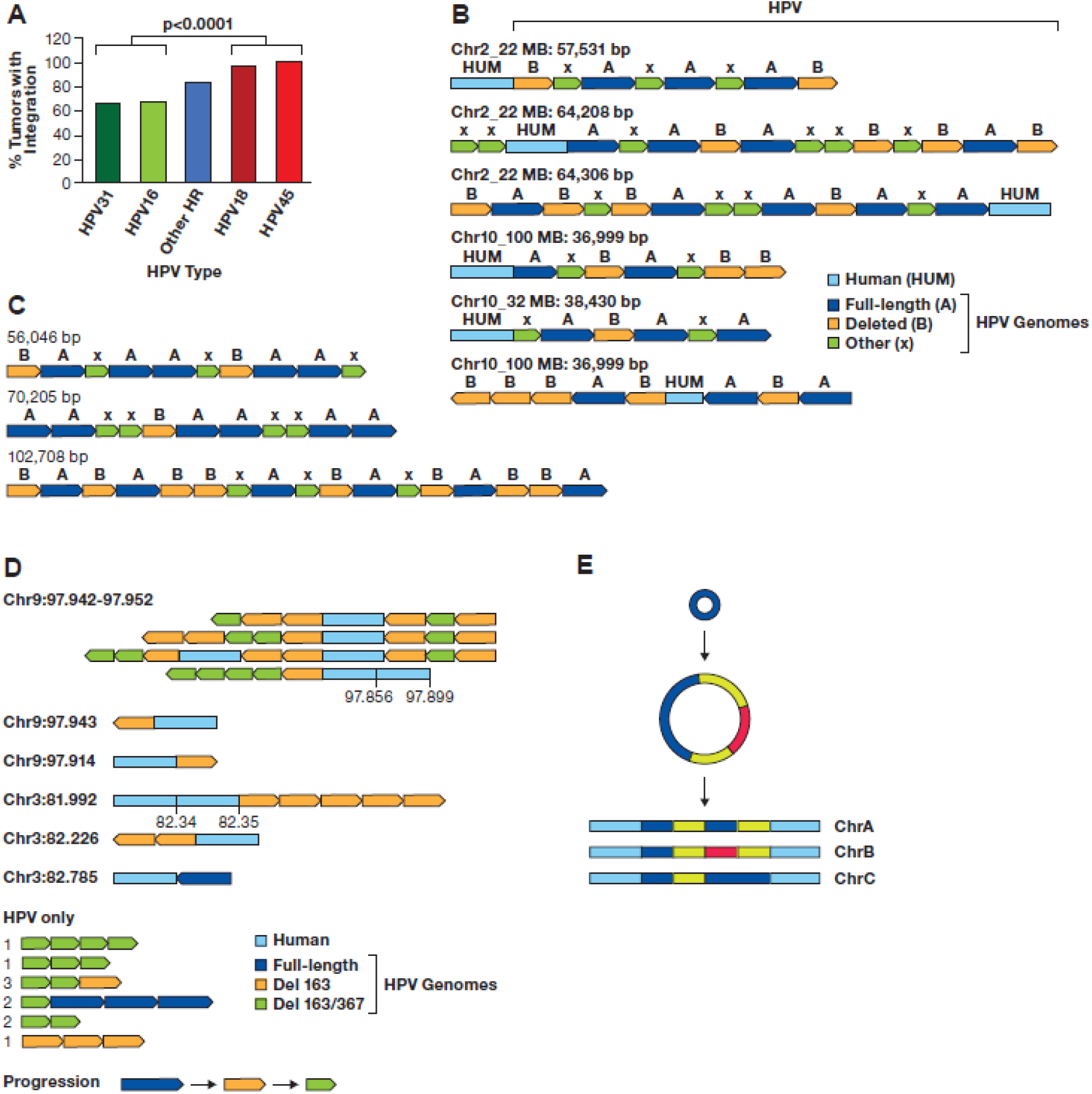
HPV multimers in cancer cell lines. **A**. The integration frequency of the predominant HPV types is shown using combined data from TCGA and Guatemalan tumors (Cancer Genome Atlas Research et al., 2017; Lou et al., 2020). The other high-risk (HR) types are combined. The number of samples is HPV31 (N=3), HPV16 (N=170), Other HR (N=43), HPV18 (N=32), and HPV45 (N=17). **B**. A diagram of contigs of long reads in CaSki cells aligning to the human genome is shown. Human DNA sequences are in light blue, the full-length HPV16 genome A in dark blue, a 6.5 kb deleted genome derived from genome A in orange, and other smaller HPV16 fragments of varying size in green (X). The arrows show the direction of the HPV genome segments. The location of the human junction and size of the contig is shown. **C**. HPV only reads from CaSki cells, key as in panel B. **D**. The location of the 163 and 367 bp deletions in the SCC152 cell line is shown, as well as their locations in the HPV16 upstream regulatory region. The 367 bp deletion removes most of the distal region, and the 163 bp deletion removes four NF1 binding sites and two YY1 sites (only one is shown) in the Intermediate Enhancer region. Shown are the binding sites for the viral E1 and E2 proteins and the transcription factors OCT1, AP1, YY1, and NF1. ORI, the origin of replication; TATA, TATA-binding site; P_97_, major promoter; P_670_, minor promoter. The locations of the coding region of the E6 and E7 genes are indicated. **E**, Contigs of SCC152 HPV16 multimers are shown, with the human/HPV junction location on the left in MB. For reads with an additional human segment, those coordinates are given above the contig. For HPV-only reads, the number of times that structure was found are shown to the left of the read diagram. A suggested progression in the evolution of the single and double deleted HPV16 genomes is shown. **F**. A model of HPV Superspreading. A monomer 7.9 kb episome can undergo aberrant replication to generate a **Multimer Episome** that may contain a combination of full-length, deleted, and rearranged HPV genomes. These structures are replicated as extrachromosomal elements but may integrate into the genome at multiple locations.

For HPV16, three classes of tumors have been described: 1) those without integration and only episomal DNA, 2) integrated only tumors, and 3) tumors with both integrated and episomal DNA (Cheung et al., 2013; Lou et al., 2020; Xu et al., 2013). To better understand the structure of HPV16 in these different tumor classes, we applied long-range, single-molecule sequencing using the Oxford Nanopore Technologies (ONT) platform. To evaluate our strategy, we first sequenced SiHa cervical cancer cells (Friedl et al., 1970), known to have a well-characterized single locus of integration on chromosome 13 (Akagi et al., 2014; Kalu et al., 2017). Using both WGS and CRISPR-targeted sequencing, we constructed a 54 kb contig containing the 7652 bp portion of the HPV16 genome and confirming the rearrangement of flanking human DNA (**Supplemental Figure 1**). Therefore, long-read and targeted sequencing can resolve the structure of HPV integration sites.

### Long-read sequencing identifies complex integrated HPV multimers

To understand the structure of complex integration events, we performed long-read sequencing of the CaSki cervical cancer cell line containing up to 800 copies of HPV16 at 30-40 chromosomal sites (Akagi et al., 2014; Baker et al., 1987). We used sheared, and unsheared DNA to achieve a range of read lengths and sequenced the libraries on the ONT platform. Reads of up to 67 kb were obtained containing HPV and human sequences, and 28 out of 35 (80%) recurrent junctions matched those seen previously using short-read WGS (**Figure 1B, Supplemental Table 1**). The HPV sequences in these reads were concatemers of:

1. a full-length genome (genome A).
2. a previously described 6.5 kb genome with a 1.4 kb deletion (genome B)(Baker et al., 1987).
3. HPV16 fragments of other sizes.

In addition, we obtained reads of up to 102 kb with only HPV16 sequences (**Figure 1C**). The HPV-only reads also contained concatemers of genomes A and B and smaller fragments. Reads aligning to the same human-HPV junction assembled into consistent contigs, but there was no identifiable pattern in the order of genomes A and B at different chromosomal locations. We subsequently constructed libraries with an ultra-high molecular weight protocol, confirmed many of these junctions and obtained HPV-human reads of up to 347 kb and HPV-only reads of up to 248 kb (**Supplemental Figure 2**). Therefore, a complex duplication and assembly process must have generated the concatemers.

We also analyzed complex integrations in a head and neck squamous cell carcinoma cell line, SCC152. This line was derived from a relapsed tumor obtained one year after the SCC090 cell line was established from the same patient (White et al., 2007). SCC090 contains 200-500 copies of HPV16 integrated at four chromosomal sites (Akagi et al., 2014). Our sequencing of SCC152 revealed nine integrated loci, supported by multiple long reads, on chromosomes 2, 3, and 9 (**Supplemental Table 2**). These integration sites were also identified in SCC090 and, therefore, retained in the relapsed tumor. Most integrated loci contained HPV16 arrays composed of 1) full-length genomes, 2) genomes containing a 163 bp deletion, and 3) genomes both the 163 bp and a 367 bp deletion (**Figure 1D, E**). However, we also observed single molecules containing HPV/human junctions on a total of 10 different chromosomes (**Supplemental Table 3**).

The 367 bp deletion in SCC152 was always seen with the 163 bp deletion, suggesting that the 163 bp deletion appeared first and the 367 bp deletion occurred on a 163 bp deletion-containing genome. Both deletions are in the URR region, with the 163 bp deletion removing part of the intermediate enhancer region and the 367 bp deletion a portion of the distal region (**Figure 1D**). As with CaSki cells, we observed a random order of the different HPV16 forms integrated into SCC152 cells at distinct genomic locations, although in SCC152, there are very few full-length genomes. The data from both CaSki and SCC152 suggest that deleted forms of HPV formed as HPV ecDNA, present as viral concatemers, before integration. We term this phenomenon of episomal amplification, episomal deletion and rearrangement, followed by integration at multiple chromosomal locations *HPV superspreading*. **Figure 1F** shows a mechanism for HPV superspreading arising from aberrant replication of episomal HPV, followed by concatemer formation, and subsequent integration. Our finding of many human/HPV junctions in common between SCC090 and SCC152, cell lines formed from separate tumors from the same patient, indicates that the integrated multimers did not occur during cell line establishment but were present in both the primary and relapsed tumors.

### A cell line with episomal HPV16 displays multimer and deleted episomes

To further understand the formation of multimer episomes and the mechanisms of HPV superspreading, we searched for cell lines with episomal HPV. The SNU-1000 cell line was established from a cervical squamous cell carcinoma isolated from a 43-year-old Korean patient and published as having episomal and integrated forms of HPV16 (Ku et al., 1997). Long-read WGS of SNU-1000 identified a 150 bp HPV fragment integrated on chromosome 11q in the intron of the *CEP126* gene. This HPV fragment contains a portion of the *E7* oncogene but cannot encode a functional E7 protein (**Figure 2A**). Read count analysis across chromosome 11 identified genes located 5’ to the integration site, such as the progesterone receptor (*PGR*) amplified 10-fold, and the *YAP1, BIRC2, BIRC3* genes, 3’ to the integration, amplified 25-fold (**Figure 2A**).

**Figure 2.**
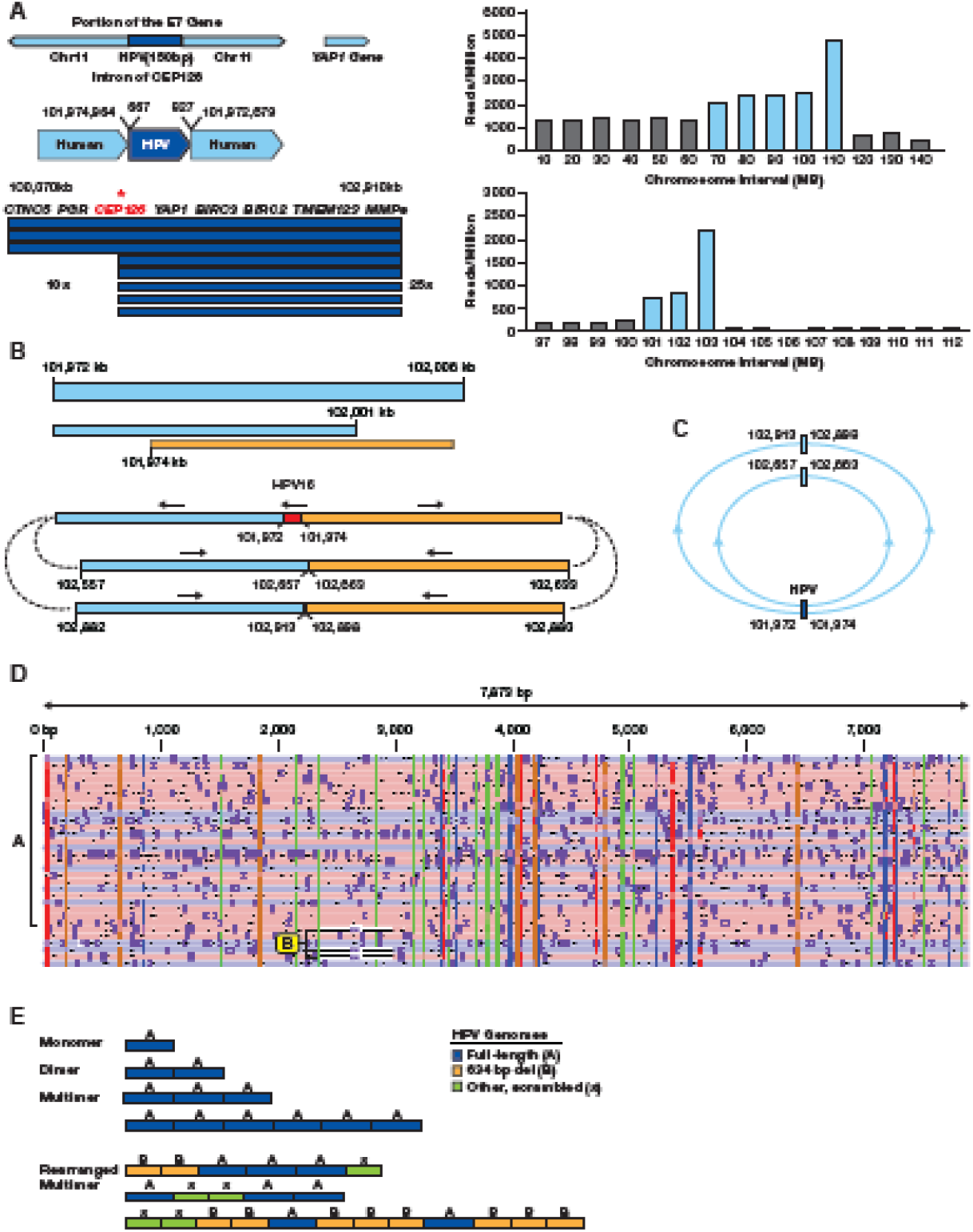
**A**. The integration locus on chromosome 11 in SNU-1000 is shown along with the fragment of HPV16 from 667-827 bp of the HPV16 genome. The arrow displays the site of integration in the *CEP126* gene and the amplified 2.9 MB region. Below are read counts normalized to reads/million total reads in 10 and 1 MB intervals across chromosome 11 and the 97-112 MB region. Most other chromosomes do not show such drastic alterations in read depth (**Supplemental Figure 3**). **B**. Long-read contigs containing the sequences flanking the integration site are shown along with the direction of the flanking human segments. Also shown are contigs of the two human-human junctions supported by multiple reads joining 102,657 to 102,663 kb and 102,913 to 102,898 kb. **C**. A model of circular structures that would be consistent with the read junctions, forming 1.4 and 1.9 MB circles. **D**. SNU-1000 WGS reads mapped to the HPV16 genome show insertions of multimers of ∼7.9 kb, as well as 634 bp deletion genomes and rearranged genomes (arrows). **E**. Representative HPV-only reads corresponding to monomers, dimers, and multimers of the full-length HPV genome and multimers containing concatemers of full-length, 634 bp deleted, and other HPV fragments are displayed.

The region distal to 11q22.2 (103-140 MB) is present at a low copy number from the WGS data, and we identified multiple long reads joining chromosome 11 at 102,657 kb to 102,663 kb and 102,898 kb to 102,913 kb (**Figure 2B**). Therefore, the chromosome 11 HPV16 integration resulted in amplifying the locus and rearranging the flanking human DNA. As we did not see chromosome 11q joined to another chromosome, this amplified DNA appears to lack a telomere. **Figure 2C** presents a model in which these sequences are in circular, extrachromosomal structures.

Sequencing of SNU-1000 DNA was performed with both standard adapter ligation onto linear DNA molecules and insertion of transposase adaptors into linear and circular molecules. In both cases, we identified large HPV16-only reads consistent with episomal DNA. We obtained transposon-tagged, HPV-only reads of 7.9 kb representing monomer episomes (**Figure 2D**). In addition, there were many reads longer than 7.9 kb containing concatemers of full-length genomes, a 634 bp deleted genome, and other truncated genomes (**Figure 2E**). Some reads contained only full-length genomes consisting of monomer episomes, dimers, and higher order concatemers. Others contained complex patterns of full-length and deleted forms indicating aberrant replication of episomal DNA. The 634 bp deletion removes a portion of the C terminus of the E1 gene and the N-terminus of E2. In addition, rare single molecules were identified in SNU-1000 joining HPV sequences to all human chromosomes except chromosome 22 (**Supplemental Table 4**). All these integration locations had support from only one read. None of these junctions were near known recurrent HPV integration sites (Tang et al., 2020), suggesting that HPV16 is randomly integrating into the human genome in subpopulations of cells during the passage of the SNU-1000 culture. Therefore, SNU-1000 displays a full spectrum of episomal monomers, multimers, deletions, and complex concatemers as extrachromosomal DNA.

### Identification of a cell line with integrated HPV18 multimers

Due to the high rate of integration of HPV18 in cancers, we were surprised to find a cell line, SNU-1245, reported to have episomal HPV18 (Ku et al., 1997). However, long-read whole-genome sequencing and targeted sequencing using computer-guided sequence selection (adaptive sampling) revealed that SNU-1245 has a single multimer integration of HPV18 on chromosome 1q32.2 (**Figure 3A**). Thus, the locus contains one full-length copy of HPV18 and three HPV18 fragments, a type 2 integration. The HPV sequence block totals 16,615 bp, and this region of chromosome 1 is amplified approximately five times. In addition, the flanking segments of human DNA are rearranged, consistent with a looping amplification mechanism, as reported previously for HPV16 and HPV18 cell lines and tumors (Akagi et al., 2014). Therefore, the structure of the SNU-1245 locus suggests that HPV18 can also undergo multimer amplification of episomal HPV DNA, followed by integration.

**Figure 3.**
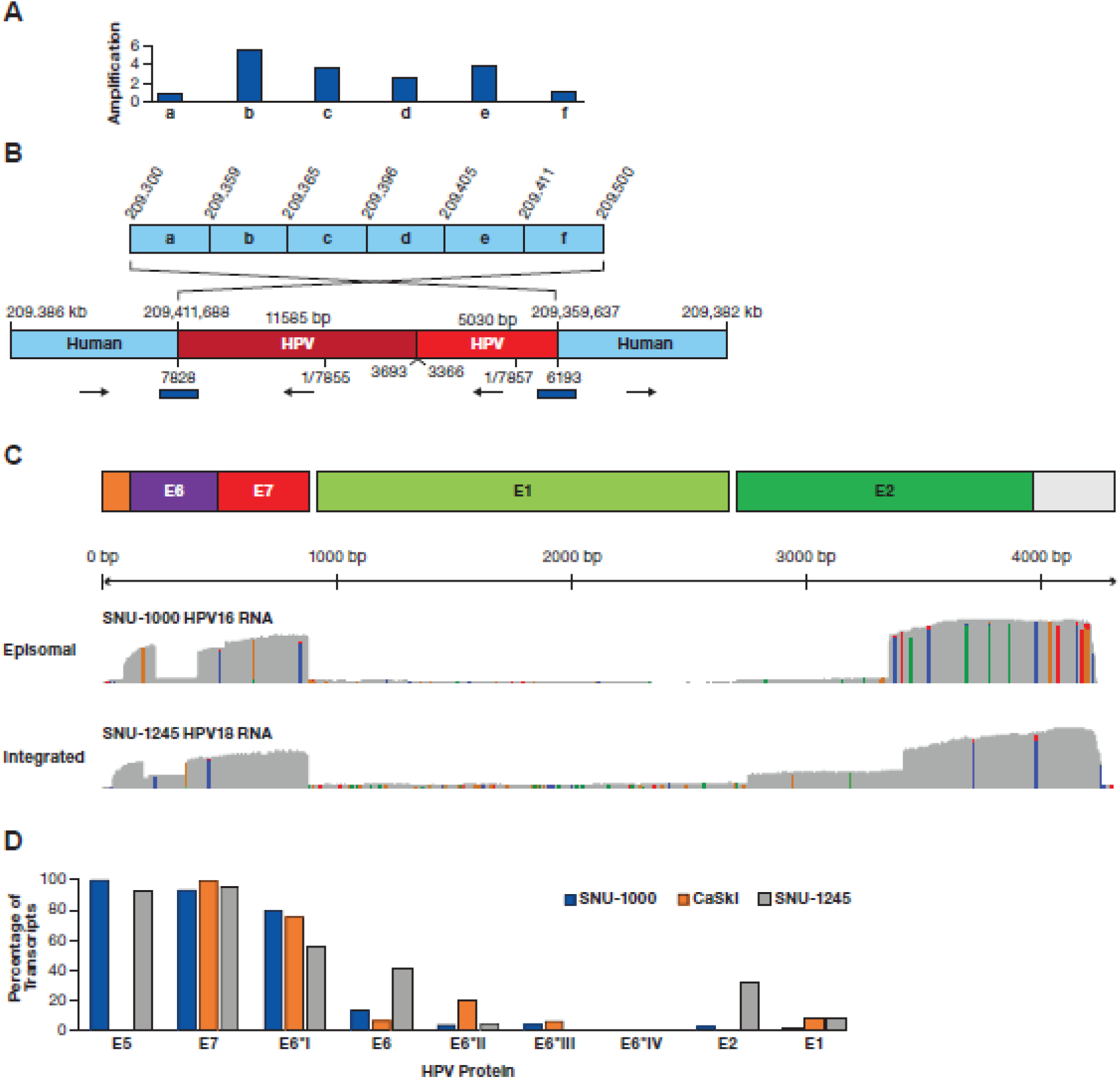
Structure of the SNU-1245 integration locus and HPV expression. **A**. A diagram of a region of chromosome 1q24.2 is shown divided into six segments (a-f). Above the diagram are copy number values normalized to segments a and f. **B**. A contig supported by multiple long reads obtained from ONT WGS, CRISPR targeted sequencing, and adaptive sampling is shown. The two human regions are rearranged, consistent with integration at 209,411,668 bp and looping back to 209,359,637 bp. Human coordinates are above the contig and HPV18 coordinates below. The total HPV contig is 16.6 kb. The blue bars represent junctions confirmed by PCR and Sanger sequencing. **C**. Plots of full-length direct cDNA sequencing are shown, displaying an abundant expression of the *E6*/*E7* gene regions with frequent splicing of E6 and deficient expression of the *E1*/*E2* gene region. **D**. The estimated level of HPV proteins is shown based on content and abundance of transcripts in **Table 1**.

### Episomal cervical cancer cells have an HPV expression profile like integrated cells

To study the expression pattern of episomal and integrated HPV, we used full-length direct cDNA sequencing to quantify the HPV16 transcripts in the SNU-1000 cell line and HPV18 transcripts in SNU-1245 (**Figure 3C**). No HPV-Human hybrid transcripts were observed in either cell line, indicating that HPV expression is within the HPV concatemers. **Table 1** shows that the most abundant HPV16 transcript (transcript B) in SNU-1000 cells encodes the spliced E6*I form of E6, E7, E4, and E5 and accounts for 78% of the transcripts. There is a very low abundance of mRNAs capable of encoding E1 or E2 in SNU-1000. In the SNU-1245 line, there is a nearly even balance between transcripts encoding the full-length E6 and E6 spliced forms. Furthermore, there is a higher level of transcripts capable of encoding E2 and E1 (transcripts 4, 5, and 6)(**Figure 3D**).

**Table 1.**
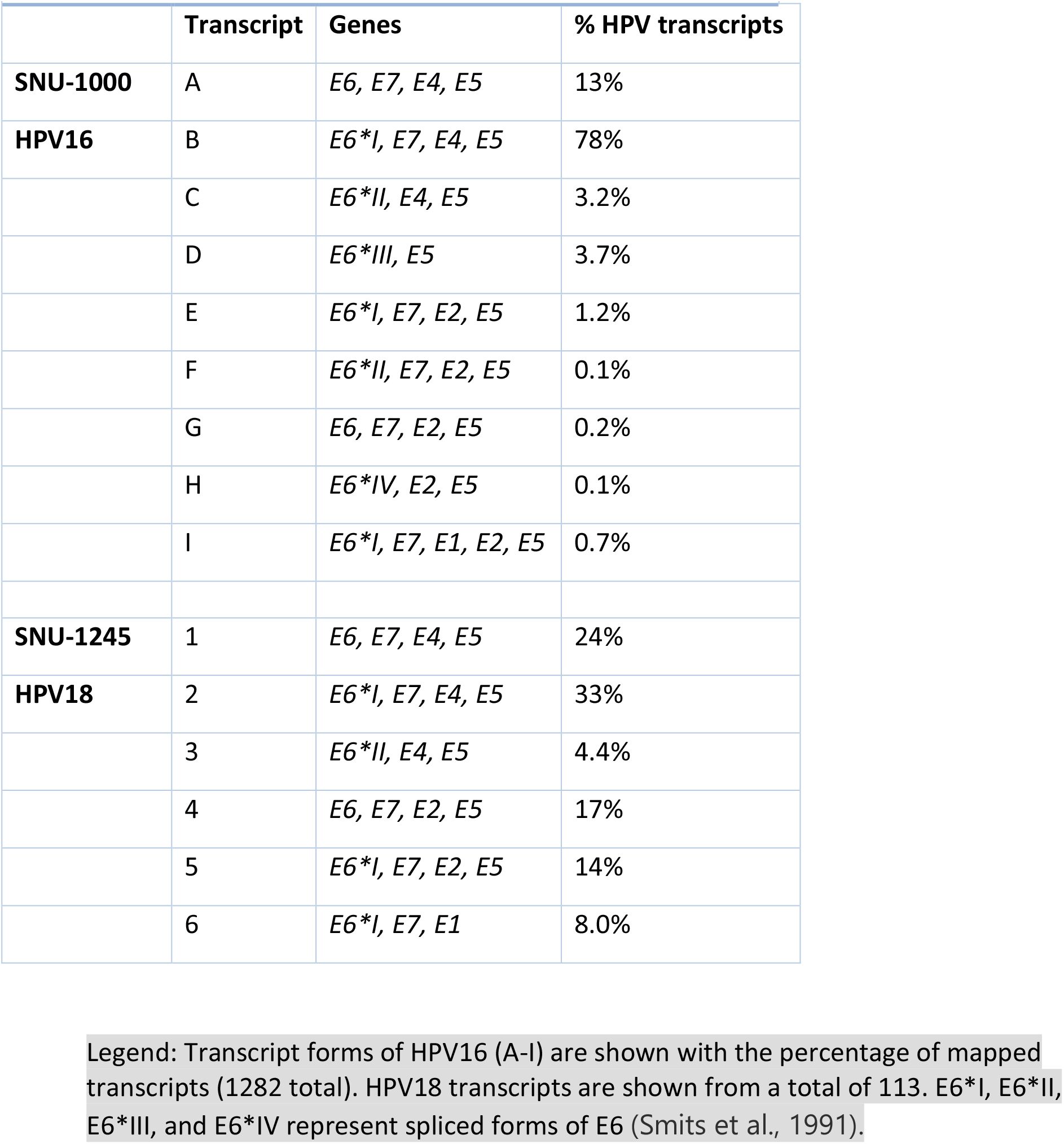
Classes of HPV16 and HPV18 transcripts.

|  | Transcript | Genes | % HPV transcripts |
| --- | --- | --- | --- |
| **SNU-1000** | A | *E6, E7, E4, E5* | 13% |
| **HPV16** | B | *E6\*I, E7, E4, E5* | 78% |
|  | C | *E6\*II, E4, E5* | 3.2% |
|  | D | *E6\*III, E5* | 3.7% |
|  | E | *E6\*I, E7, E2, E5* | 1.2% |
|  | F | *E6\*II, E7, E2, E5* | 0.1% |
|  | G | *E6, E7, E2, E5* | 0.2% |
|  | H | *E6\*IV, E2, E5* | 0.1% |
|  | I | *E6\*I, E7, E1, E2, E5* | 0.7% |
| **SNU-1245** | 1 | *E6, E7, E4, E5* | 24% |
| **HPV18** | 2 | *E6\*I, E7, E4, E5* | 33% |
|  | 3 | *E6\*II, E4, E5* | 4.4% |
|  | 4 | *E6, E7, E2, E5* | 17% |
|  | 5 | *E6\*I, E7, E2, E5* | 14% |
|  | 6 | *E6\*I, E7, E1* | 8.0% |

Legend: Transcript forms of HPV16 (A-I) are shown with the percentage of mapped transcripts (1282 total). HPV18 transcripts are shown from a total of 113. E6\*I, E6\*II, E6\*III, and E6\*IV represent spliced forms of E6 (Smits et al., 1991).

### Multimer Episomes are present in cervical tumors

To analyze the structure of HPV16 episomes in cervical tumors, we performed ONT tagmentation sequencing on tumors previously classified as episomal (EP) only or episomal and integrated (EP/INT)(Lou et al., 2020). The age range, *PIK3CA* mutation status, histology, HPV type, and HPV16 sublineage are shown in **Figure 4A**. From 28 HPV16 EP tumors, we obtained sufficient HPV16-containing reads to examine the structure of the 20 episomal DNAs. None of these tumors had HPV/human junction reads, consistent with their status as EP. For eight of these tumors, the longest reads began and ended at nearly the same position on the 7906 bp viral genome and had no insertions, deletions, or rearrangements (**Supplemental Table 5**). This result is consistent with these molecules representing circular episomes, tagged in random positions by the transposon (**Figure 4B**). These tumors and those with multiple reads all less than 7906 bp were classified as monomer-only tumors. Therefore, approximately one-half of EP tumors retain a monomer episome.

**Figure 4.**
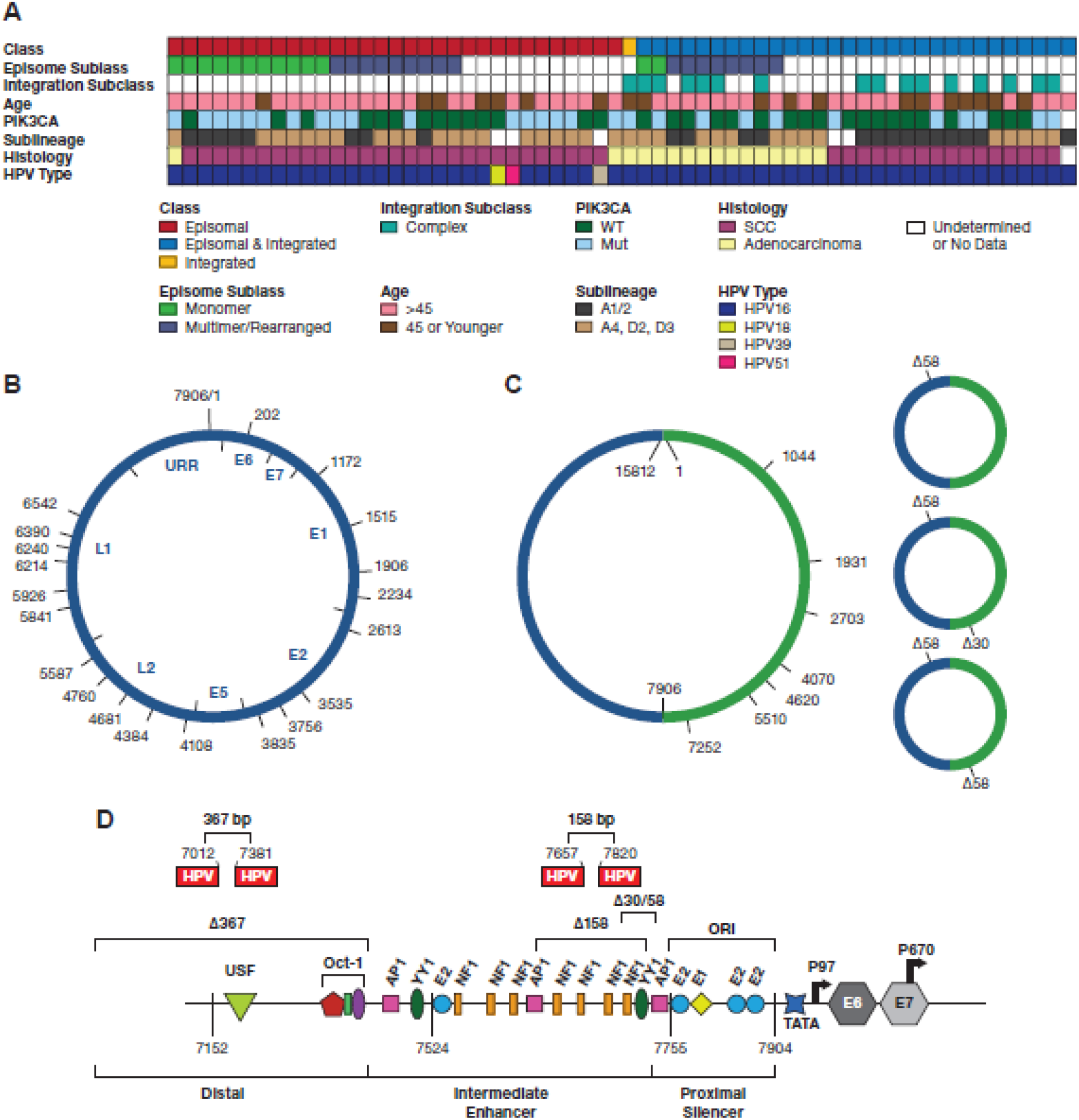
**A**. Diagram displaying data from 62 tumors. The integration class and subclass, episomal subclass, age range, *PIK3CA* mutation status, HPV type and HPV16 sublineage, and histology are shown. SCC, squamous cell carcinoma; WT, wild type; Int, integration. Blanks represent samples not able to be classified. **B**. Diagram of the 7906 bp HPV16 genome DNA, displaying the start site of DNA sequence reads from 13 tumors containing a complete or nearly complete genome sequence. The position of the HPV16 genes and URR are shown inside the diagram. **C**, A diagram of HPV16 dimers from three tumors showing the start position of reads with complete or nearly complete copies of two tandem HPV16 genomes (**Supplemental Table 6**). **D**, the location of 30 and 58 bp deletions in the URR occurring in dimer reads from tumor T393.

A total of nine other EP tumors had either HPV16 DNA reads larger than 7.9 kb, representing multimer episomes, or had rearranged HPV-HPV junctions. Three of these tumors had sequences of 15.8 kb with two complete copies of the HPV16 genome, representing HPV dimers (**Figure 4C**). As with the monomers, these reads start and stop at nearly the same position on the genome. One tumor, T393, had dimers with both 30 and 58 bp deletions in the URR (**Figure 1D**). In addition, dimer reads with one or two copies of Δ58 or one copy of Δ30 and Δ58 were observed (**Figure 4D**). The Δ30 and Δ58 deletions are in overlapping regions of the URR and delete an NF1 binding site and two YY1 binding sites. (**Figure 1D**). These two YY1 binding sites have previously been found deleted in HPV16 isolates from cervical cancer, leading to elevated activity of the P_97_ promoter (May et al., 1994). Therefore, as seen in the SNU-1000 cell line, deletions can propagate in episomes replicating in cervical tumors and give rise to multiple aberrant structures.

Six other EP tumors had episomal DNA with rearrangement of HPV16 sequences (**Figure 5A**). Interestingly, many of these tumors have a breakpoint inside the *E1* (914-2666 bp) or E2 genes (2756-3853). These rearrangements could separate the E1 and *E2* open reading frames from the P_97_ promoter and lead to decreased expression of *E1* and *E2* or disrupt their reading frame. Most of these HPV16 rearrangement junctions were observed in our previously published HPV capture and Ion Torrent sequencing (Lou et al., 2020), performed on the same DNA samples (**Supplemental Figure 4**). This data clearly shows that rearrangement of episomal DNA is a common feature of HPV16-driven cervical tumors. In addition, transcriptome analysis of representative EP-only tumors revealed that E6/E7 transcripts are predominant and that nearly all splice inside the E6 gene (**Supplemental Figure 5**).

**Figure 5.**
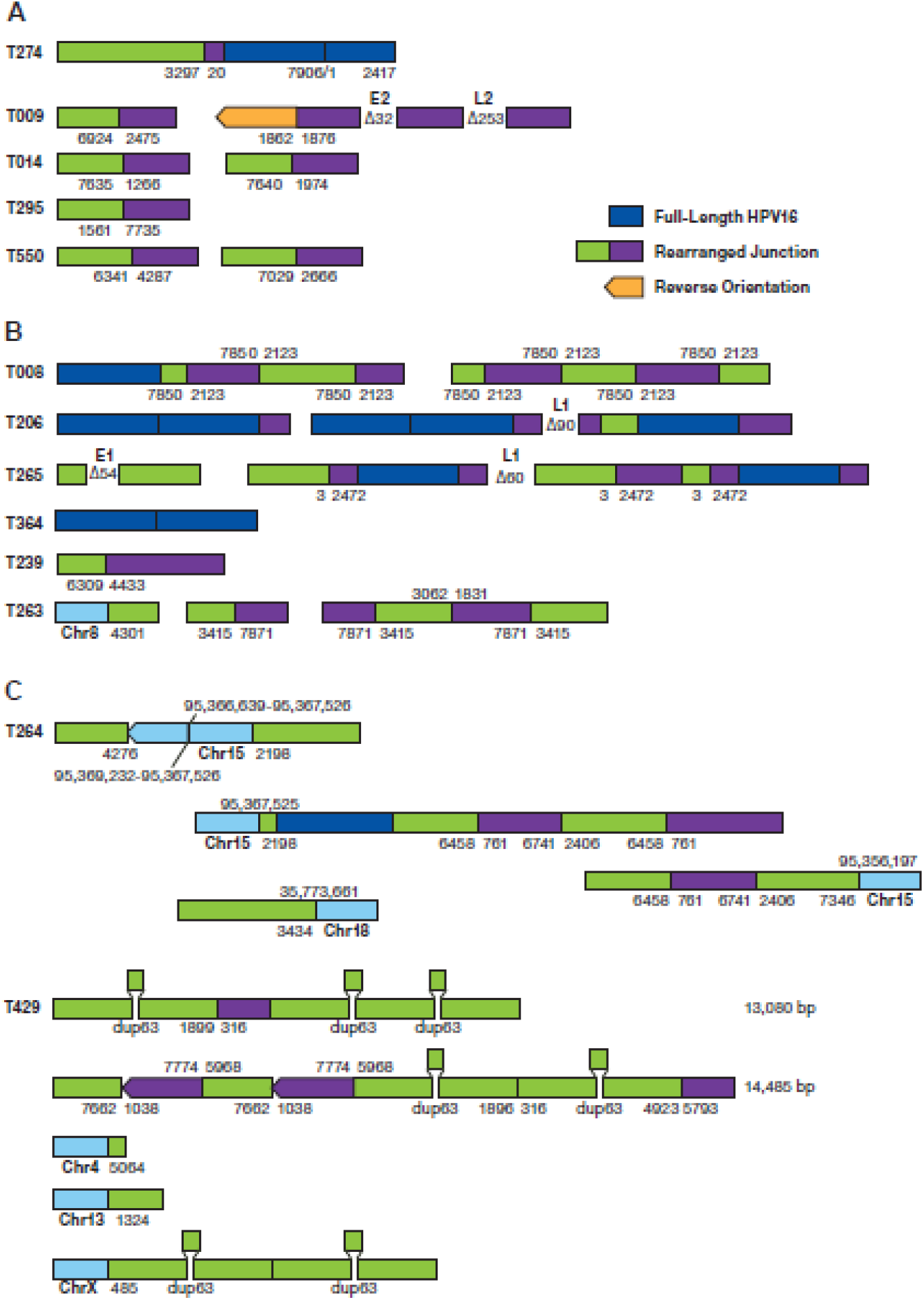
Structure of HPV sequences in Episomal and Integrated tumors. **A**. The structure of rearranged HPV-only reads in EP tumors is shown. The numbers represent the position of the HPV16 genome at the site of the breakpoint. Deletions are displayed between segments. Unless shown, all HPV sequences are arranged in the same orientation. **B**. Structure of HPV reads in EP/INT tumors. Regions of human DNA are shown in light blue. **C**. The structure of HPV reads in tumor T429 is shown. HPV16 sequence breakpoints are shown joining chromosomes 4, 13, and X to HPV sequences. The position of a 63 bp duplication is shown. The top 2 diagrams represent independent sequence reads of 13,085 and 14,485 bp.

Long-read WGS of EP/INT tumors gave a more complex picture. Despite having only 0.5x genome coverage, we identified the same HPV-human breakpoint in 13/27 tumors found by HPV capture and short-read sequencing (Lou et al., 2020). Nearly all tumors for which the integration was not confirmed by ONT sequencing yielded less than 10 HPV-containing reads. Many EP/INT tumors have complex rearrangements of HPV-only sequences (**Figure 5B**). Several tumors have 50-100 bp deletions internal to the HPV sequence or have multiple rearranged junctions. As with EP tumors, many HPV junctions are within the 865-3875 region of the HPV16 genome containing the *E1* and *E2* genes.

Tumor 429, an EP/INT tumor with integrated DNA in multiple chromosomes, contains a recurrent 63 bp duplication in the E1 gene. The deletion was seen twice each in single HPV-only molecules of 13 and 14 kb (**Figure 5C**). In addition, the dup63 mutation was seen twice in a read anchored to chromosome X. Therefore; this is a case of superspreading; aberrant replication of episomal DNA that integrates into multiple chromosomal locations in a cervical tumor.

## DISCUSSION

Human papillomaviruses are among the most oncogenic human cancer viruses, and HPV16 is the most carcinogenic type of HPV (Mirabello et al., 2018; Mirabello et al., 2016; Schiffman et al., 2016). While integrating viral DNA into the human genome is an important mechanism of HPV carcinogenesis, transformation without HPV integration is less well understood. Our data employing long-read and single-molecule sequencing of cell lines and cervical tumors reveal new aspects of HPV oncogenesis. By carrying out long-read WGS on well-characterized cell lines (SiHa, CaSki, SCC152), we established that the method reliably identifies HPV sequences and the structure of integrated loci. Using HPV-containing reads of up to 347 kb, we show that many of the integrated loci in the CaSki cell line are composed of complex strings of full-length genomes, a recurrent 6.4 kb truncated genome and other smaller fragments. The finding of concatemers of similar structure and complexity integrated into multiple regions of the genome supports a model where the mixed concatemers of HPV genomes were generated as extrachromosomal DNA and subsequently inserted into the human genome. This model is further supported by the head and neck cancer cell line, SCC152, which has unique HPV16 forms with Δ163 bp and Δ163/Δ367 bp deletions. In addition, SCC152 has complex concatemers integrated at different sites on chromosomes 3 and 9. We term this phenomenon of extrachromosomal HPV rearrangement and amplification, followed by integration, **HPV Superspreading**.

To support the superspreading model, we analyzed a unique cell line, SNU-1000, that stably replicates HPV16 episomal DNA (Ku et al., 1997). Established from a Korean squamous cell carcinoma, SNU-1000 contains only a 150 bp fragment of the HPV16 E7 gene integrated on chromosome 11q22.1. Interestingly, this integration site is within a 2.9 MB amplified region containing the *YAP1, BIRC2*, and *BIRC3* genes. Thus, nearly all the remaining HPV16 sequences in SNU-1000 are in molecules devoid of human DNA and appear extrachromosomal. To our knowledge, this is the first time an oncogene amplification has been found at a locus with only an incomplete copy of the *E6* or *E7* genes, representing a new mechanism of HPV carcinogenesis.

Using a transposon-based approach that can directly sequence linear and circular DNA, we recovered multiple reads from SNU-1000 that begin and end at the same or nearly the same position on the circular HPV16 genome. These reads are almost certainly derived from intact monomer episomes. We also identified reads that represent intact dimers and multimers of full-length HPV16 genomes, indicating that replication of the HPV16 monomer can lead to intact multimer genomes. Multimeric forms of HPV16 and HPV18 have been observed in cell lines and transfected cells, indicating that this may be common in HPV pre-cancers and cancers (Choo et al., 1989; Hall et al., 1997; Orav et al., 2013). We further analyzed HPV16 sequences in SNU-1000 and identified a common, recurrent 634 bp deletion, removing portions of the *E1* and *E2* genes. This deletion is present in about 15% of viral genomes and occurs exclusively in large multimer episomal structures. These multimer episomes are composed of concatemers of full-length HPV16 genomes, Δ634, and other rearranged genomes.

Analysis of direct, full-length cDNA from SNU-1000 demonstrates that the predominant HPV transcript in these cells encodes the E6*I, E7, E4, and E5 proteins and low amounts of E1 and E2 encoding transcripts. Thus, in the absence of integration, rearranged and deleted HPV16 multimers appear to have resulted in the upregulation of E6 and E7 and downregulation of E1 and E2, a similar profile as tumors with integrated HPV. In addition, equal or higher levels of E6 and E7 in episomal HPV16-positive cervical cancers have been observed (Cheung et al., 2013; Hafner et al., 2008). Therefore, one mechanism of HPV carcinogenesis, in the absence of integration, is the episomal deletion of *E1* or *E2* (**Figure 6**). As *E1* and *E2* are frequently deleted or inactivated in integrated HPV, extrachromosomal alteration of these genes may be selected for upregulating E6 and E7 expression.

**Figure 6.**
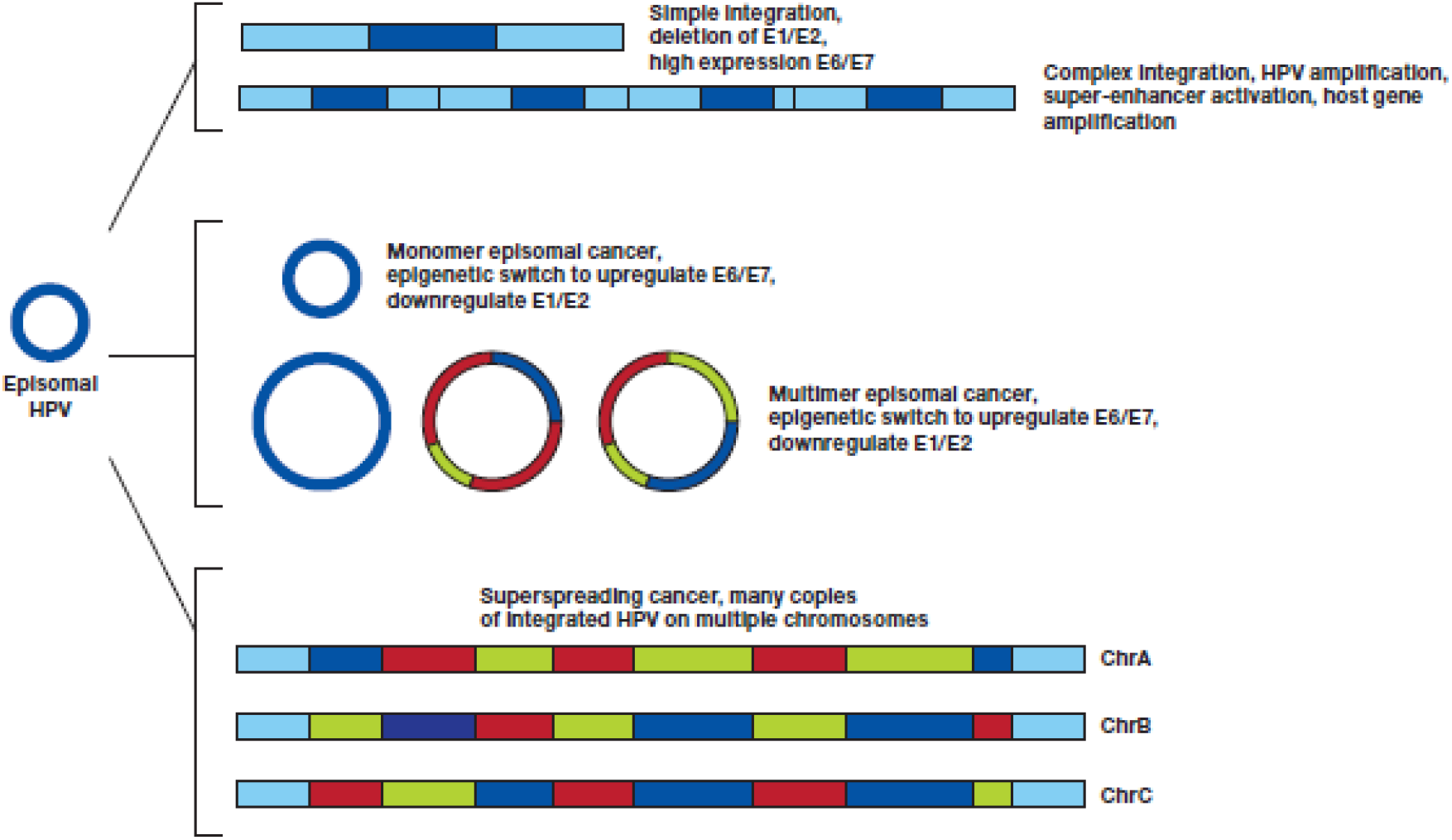
Mechanisms of HPV carcinogenesis. An HPV episome can undergo a **simple integration**, deleting the *E1* and *E2* genes, inhibiting DNA synthesis at the HPV origin of replication (ORI) and upregulating E6 and E7. If during integration E1 and E2 are not suppressed, a local amplification can case a **complex integration** leading to many copies of HPV and flanking DNA, resulting in higher expression of E6 and E7, formation of a super-enhancer activating flanking genes (Warburton et al., 2018), and potentially amplifying a carcinogenic host gene. **Monomer episomal cancer** occurs when the genome remains an episome but undergoes an epigenetic switch to upregulate E6/E7. Aberrant episome replication can lead to **Multimer episomal cancer** in which deletions and rearrangements in the E1/E2 genes or URR lead to high E6/E7 and low E1/E2 expression. Insertion of multimer episomes into multiple genomic locations give rise to HPV **superspreading cancer**.

To better understand the mechanisms of carcinogenesis in tumors without HPV16 integration, we studied samples previously characterized as episomal-only. Interestingly, nearly one-half of these tumors have intact monomer episomes. Therefore, HPV16 is capable of causing cancer without integration or rearrangement of episomal DNA. Monomer-only tumors were mostly SCC and included HPV16 A1, D2, and D3 sublineages. However, RNAseq data shows a transcription pattern dominated by the expression of mRNAs containing the E6 and E7 genes. This finding indicates that HPV episomes have undergone an epigenetic switch to increase the expression of the E6 and E7 oncogenes. Exploring the molecular basis of this alteration in viral gene regulation may provide further insight into HPV-driven cancers.

We identified a subset of episomal-only tumors with rearranged and deleted episomal HPV. Furthermore, these deletions and rearrangements nearly always have an HPV-HPV breakpoint in the *E1* or *E2* genes, supporting a model where episomal disruption of E1/E2 can contribute to cancer. It is interesting to speculate that as multimer episomes expand, they gain more replication origins, increasing the rate of further expansion and deletion. Therefore, HPV16 can cause cervical cancer without integration via a combination of episome deletion and rearrangement and favoring E6/E7 oncogene expression.

Integration is not part of the typical lifecycle of HPV and results in a dead-end for the virus. Integrated forms of HPV nearly always retain the URR, and the origin of replication (ORI) is in the URR. As the E1 and E2 proteins promote replication at the ORI the presence of both episomal and integrated HPV presents a situation where E1 and E2 could promote unscheduled replication at the ORI in integrated HPV. Orav et al. and Peter et al. observed DNA synthesis at an integrated ORIs in transfected cells and proposed that this ‘re-replication’ leads to local amplification of HPV and human DNA, the formation of viral/human DNA-containing ecDNA, and chromosome translocations (Orav et al., 2015; Peter et al., 2010). Akagi et al. used WGS to characterize HPV integration sites with local amplification. They proposed a looping model whereby HPV integration itself causes the amplification of viral and flanking human DNA (Akagi et al., 2014). Finally, using computational methods, Kim et al. have demonstrated that circular ecDNA is frequent in solid tumors and 12% of cervical tumors (Kim et al., 2020). We reanalyzed the Kim et al. data and found that 38% of integrated HPV16 cervical tumors are predicted to have circular ecDNA (**Supplemental Table 7**). While this work was in preparation, Pang et al. used short-read sequencing and computational analyses to identify ecDNA, containing human and HPV sequences, in over one-third of HPV+ HNSCC tumors (Pang et al., 2021). However, our sequencing data does not directly address these post-integration events. We do show that rearrangement and amplification of HPV can occur before integration. Deleting *E1* and *E2* in episomal DNA, and epigenetic silencing, may permit an integration locus with an ORI to remain relatively stable in the presence of episomal DNA.

There are several methods to identify the presence of HPV integration, either using DNA or RNA-based analyses. We used the presence or absence of an HPV/human DNA junction to determine integration status. In this context, we classified tumors without a detected integration and an intact viral genome as episomal-only. As all classical type 1 HPV integrations involve a deletion in the *E1*/*E2* region, we used the sequence of overlapping amplicons spanning the entire HPV genome to detect deletions in HPV coverage and classify tumors with complete deletions in the *E1*/*E2* region as integrated-only. The classification of tumors as episomal and integrated is more problematic. As we and others have shown, tumor cell lines such as CaSki and SCC152 can have full-length copies of HPV16 integrated into the genome (type 2 integration) and therefore be confused as EP/INT. SNU-1245 was classified as EP/INT based on an intact *E2* gene; however, it contains a full-length copy of HPV18 integrated into the genome and has no episomal DNA. Morgan et al. argue that, at least for HNSCC, the EP/INT category is mischaracterized and that many of these tumors have ecDNA containing viral and human sequences (Morgan et al., 2017). Our data show that EP/INT tumors can have monomer, multimer, or rearranged HPV-only sequences. And we found at least one tumor with an integrated HPV multimer. However, without very deep, long-read sequencing, it is not possible to determine if these sequences are episomal or integrated multimers or if hybrid HPV/human ecDNA is present.

This study has several limitations, including that we based some of the analyses on a limited set of cell lines and have only one cell line with episomal HPV16. There have been few high-risk HPV-containing cell lines with episomal DNA described, and they tend to be unstable, losing episomal DNA over time (Jeon et al., 1995). In addition, SNU-1000 may be unique in having amplified *YAP1*, a potent cervical cancer oncogene (Lorenzetto et al., 2014). In mouse models, high *Yap1* gene expression gave rise to cervical tumors in the absence of HPV oncogenes (He et al., 2019). Therefore, YAP1 overexpression could aid in the generation of additional episomal HPV models. Due to limitations in DNA quantity and quality, we obtained relatively low coverage WGS of tumors. Therefore, we have not detected all HPV-containing species, and our tumor classifications are incomplete. However, to our knowledge, this is the first study to apply long-read sequencing to the study of episomal HPV in cervical tumors.

In conclusion, we find that multimers of the HPV genome are generated in cervical tumors replicating as extrachromosomal episomes. This HPV replication is commonly associated with deletion and rearrangement of the HPV genome. We confirm by DNA and RNA sequencing that a subset of HPV16-containing tumors has only episomal viral DNA. About half of episomal tumors have intact monomer episomes and an expression pattern dominated by E6 and E7 expression. Another subset of episomal tumors has rearranged episomes, often deleting the *E1* and *E2* genes. Our data support a model of HPV-only structures replicating as ecDNA, accumulating rearrangements leading to the integration of rearranged HPV multimers in the human genome (**Figure 6**). This process parallels the well-described amplification of oncogenes and drug resistance genes as ecDNA or double minute chromosomes that integrate as homogenously stained regions (HSRs) (Carroll et al., 1988). Further study of HPV extrachromosomal amplification and integration may provide insight into gene amplification and ecDNA formation across cancer types.

## STAR METHODS

### Patients and Informed consent

This study was conducted at the Instituto de Cancerología (INCAN) in Guatemala City and the Hospital Central Universitario “Dr. Antonio M Pineda,” Barquisimeto, Lara State, Venezuela. INCAN is the largest cancer hospital in Guatemala and the Hospital Central Universitario is a reference center for obstetrical tumors. The Research Ethical Committees of both institutions approved the protocol, and the study was declared the study exempt from institutional review board (IRB) approval by the NIH Office of Human Studies Research. Women gave written informed consent. Women over the age of 18 referred for a diagnostic biopsy to confirm cervical cancer were recruited into the study. There were no other inclusion criteria, and the only exclusion was for women unable or unwilling to provide informed consent. The subjects were not consecutive patients, and we estimate that we captured 10-20% of women diagnosed with breast cancer at these centers.

### Cell culture, DNA and RNA extraction

The cervical cancer cell lines were obtained from American Type Culture Collection (ATCC) and the Korean Cell-Line Bank and Cancer Research Center, Seoul National University (Ku et al., 1997). Cells were cultured in EMEM or RPMI-640 media with 10% Fetal Bovine Serum 1% Pen Strep (10,000 units/mL of penicillin, 10,000 µg/mL of streptomycin, and 25 µg/mL of Amphotericin B) until 70-80% confluent. Cells were washed with 10 mL PBS and harvested using 2 mL trypsin per T-75 flask. DNA was extracted using a Gentra Puregene kit from Qiagen or Circulomics Nanobind HMW DNA extraction kit. Cell line RNA was prepared from 30 million cells using Trizol (ThermoFisher) and Poly-A+ RNA purified by DYNAL Dynabeads (Invitrogen). Tumor DNA and RNA were simultaneously purified from 1cm of tumor tissue stored in RNALater using the DNeasy Blood & Tissue Kit or RNeasy Mini kit. DNA was quantitated by Nanodrop (Thermo Scientific) and Qubit (Thermo Scientific) and stored at 4°C; RNA was quantitated by Qubit and stored at −80°C.

### Long-read DNA sequencing

For ligation sequencing of cell line DNA, 1 ug was either sheared to 8-20 kb with a G-tube (Covaris) or used unsheared with the LSK-109 kit according to instructions. Targeted sequencing of HPV16 was carried out in CaSki and SiHa cells using CRISPR probes to HPV16 probes designed using CHOPCHOP (chopchop.cbu.uib.no) and TP53 probes described previously (Gilpatrick et al., 2020) with the Cas9 targeted sequencing protocol (Oxford Nanopore). For transposase sequencing of cell lines 1 ug was used with the Rapid Sequencing kit (SQK-RAD004, Oxford Nanopore). CaSki and SNU-1000 DNA was also prepared from 6 million cells using the Ultra-Long protocol of Circulomics and the Ultra-Long DNA Sequencing kit (SQK-ULK001, Oxford Nanopore) both with and without adaptive sampling using a combined human HG38 genome and high-risk HPV type FASTA file, and a BED file selecting for cancer genes, integration loci and HPV.

Tumor DNAs (0.3 ug) were sequenced using the Rapid Barcoding kit (SQK-RBK004, Oxford Nanopore) in groups of 6 tumor DNAs. For all DNA samples, A total of 30-50 fm of DNA libraries were loaded onto MinIon R9.4 flow cells (Oxford Nanopore). In some runs, flow cells were flushed at 24 and 48 hrs and reloaded.

### Full-length RNA sequencing

Cell line RNA (500ng Poly-A+) was sequencing using the Direct RNA sequencing kit (SQK-RNA002, Oxford Nanopore). Tumor RNA (50ng total RNA) was sequenced with the PCR-cDNA Barcoding kit (SQK-PCB109, Oxford Nanopore) in groups of six samples. RNA libraries were loaded onto MinIon R9.4 flow cells on a GridIon instrument (Oxford Nanopore).

### PCR and Sanger sequencing validation

The Primer3 online program (Untergasser et al., 2012) was used to design primers for PCR amplification. Overlapping primers were designed to span the entire 7.9 kb HPV16 and HPV18 genomes. Cell line DNA was amplified using a long-range PCR kit from New England BioLabs. The samples were run on a 1.5% agarose gel at 100V and imaged with a BioRad ChemiDoc Image System. Sanger sequencing was performed using an Applied Biosystems® 3500xL Genetic Analyzer from Thermofisher Scientific.

### Bioinformatics and statistics

The fastq files from sequencing runs were aligned to the human genome (HG38) or HPV genomes using EPI2ME server (https://epi2me.nanoporetech.com) using the Fastq Human Alignment GRCh38 or Fastq Custom Alignment app with a HPV16 or HPV18 or 13 high-risk HPV type fasta file. Read alignment data for HG38 and HPV were merged with in Excel or in Filemaker (Claris) and merged with read length and adaptive sampling decision data, where appropriate. Reads of interest were manually extracted and mapped using BLAT (Kent, 2002) (https://genome.ucsc.edu) or BLAST.

BAM files produced by EPI2ME were merged and indexed using the BamTools Merge and SAMtools Index tools in the Cancer Genomics Cloud (CGC)(Lau et al., 2017). A ONT WGS Data Processing pipeline, based on a Broad Institute pipeline, was also run on CGC to align human and HPV reads, produce BAM files. BAM files were viewed in the Integrated Genome Viewer (Thorvaldsdottir et al., 2013).

DNA reads were also analyzed from fast5 raw data by base-calling using Guppy/4.5.4 (https://nanoporetech.com/). Modified base-calling was performed using Megalodon/2.3.3 https://github.com/nanoporetech/megalodon. Structural variation calling was carried out with the Nanopore pipeline-structural-variation with modifications. The entire workflow is available at https://github.com/NCI-CGR/Nanopore_DNA-seq. RNA reads were processed using the fast5 raw data and base-called using BINITO /v0.3.7 and aligned to the HG38 genome using Minimap2/2.17. Isoforms were detected and quantified using Stringtie2/2.1.5 (Kovaka et al., 2019) and Freddie (https://www.biorxiv.org/content/10.1101/2021.01.20.427493v1.full) (https://github.com/vpc-ccg/freddie). The Freddie program calls the Gurobi package (www.gurobi.com) to solve optimization problems. The entire workflow is available at https://github.com/NCI-CGR/Nanopore_RNA-seq.

Statistical analysis was performed in GraphPad.

## Acknowledgements

Thanks to Lineth Boror, Ester Avila and Patricia Zaid for sample collection. The authors acknowledge the research contributions of the Cancer Genomics Research Laboratory for their expertise, execution, and support of this research in the areas of project planning, wet laboratory processing of specimens, and bioinformatics analysis of generated data. This project has been funded in whole or in part with Federal funds from the National Cancer Institute, National Institutes of Health, under NCI Contract No. 75N910D00024. The content of this publication does not necessarily reflect the views or policies of the Department of Health and Human Services, nor does mention of trade names, commercial products, or organizations imply endorsement by the U.S. Government. We are grateful for the use of the NIH Helix Biowulf computing facility. The Seven Bridges Cancer Research Data Commons Cloud Resource has been funded in whole or in part with Federal funds from the National Cancer Institute, National Institutes of Health, Contract No. HHSN261201400008C and ID/IQ Agreement No. 17×146 under Contract No. HHSN261201500003I and 75N91019D00024.

## Author contributions section

Declaration of interests

“The authors declare no competing interests.”

## Notes

### Competing Interest Statement

The authors have declared no competing interest.

