## Supplemental_Figures for "Extrachromosomal Amplification of Human Papillomavirus Episomes as a Mechanism of Cervical Carcinogenesis"

### Slide 1
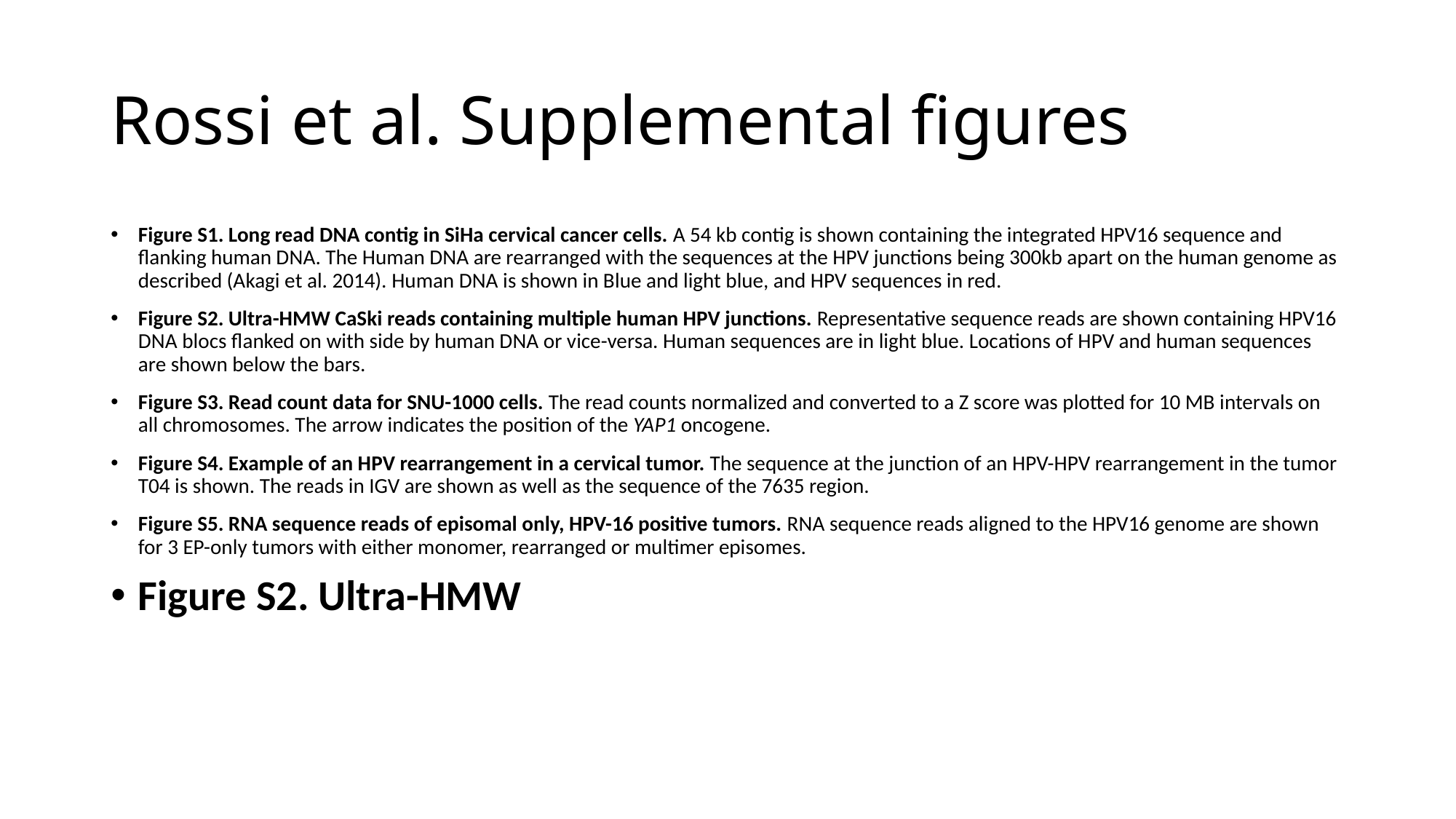

#### Rossi et al. Supplemental figures
Figure S1. Long read DNA contig in SiHa cervical cancer cells. A 54 kb contig is shown containing the integrated HPV16 sequence and flanking human DNA. The Human DNA are rearranged with the sequences at the HPV junctions being 300kb apart on the human genome as described (Akagi et al. 2014). Human DNA is shown in Blue and light blue, and HPV sequences in red.
Figure S2. Ultra-HMW CaSki reads containing multiple human HPV junctions. Representative sequence reads are shown containing HPV16 DNA blocs flanked on with side by human DNA or vice-versa. Human sequences are in light blue. Locations of HPV and human sequences are shown below the bars.
Figure S3. Read count data for SNU-1000 cells. The read counts normalized and converted to a Z score was plotted for 10 MB intervals on all chromosomes. The arrow indicates the position of the YAP1 oncogene.
Figure S4. Example of an HPV rearrangement in a cervical tumor. The sequence at the junction of an HPV-HPV rearrangement in the tumor T04 is shown. The reads in IGV are shown as well as the sequence of the 7635 region.
Figure S5. RNA sequence reads of episomal only, HPV-16 positive tumors. RNA sequence reads aligned to the HPV16 genome are shown for 3 EP-only tumors with either monomer, rearranged or multimer episomes.
Figure S2. Ultra-HMW

### Slide 2
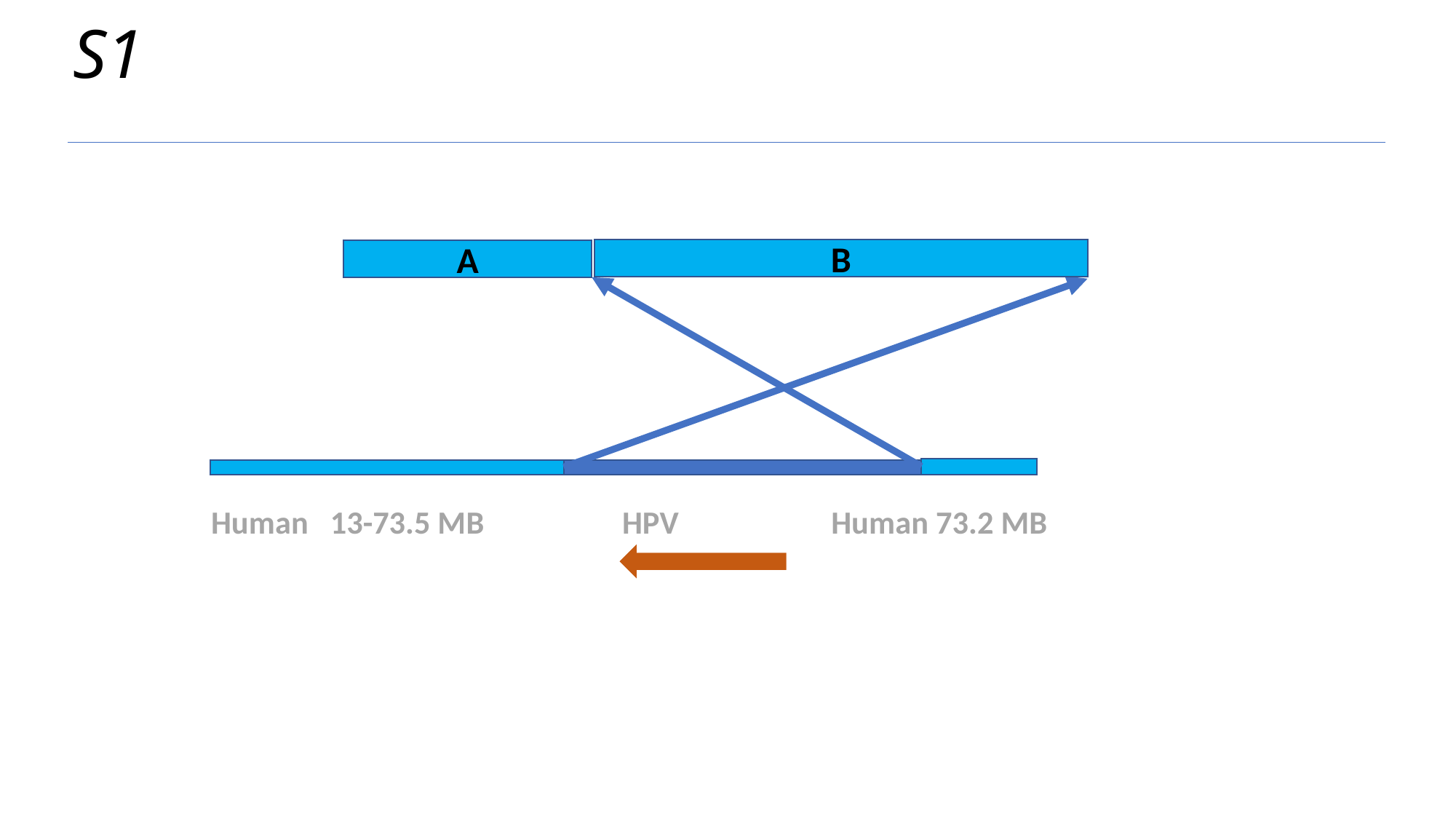

S1
B
A
Human 13-73.5 MB HPV Human 73.2 MB

### Slide 3
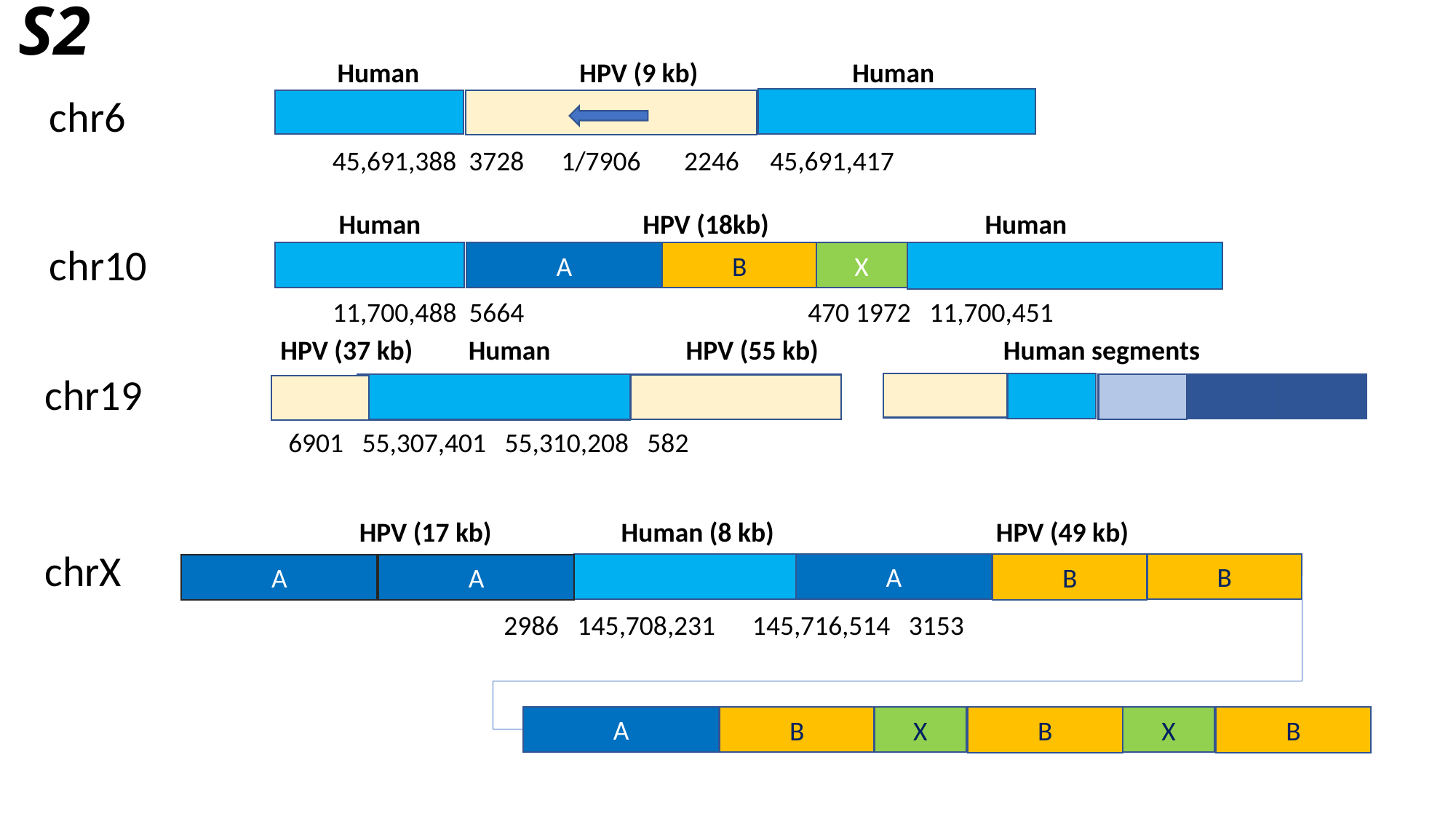

# S2
Human HPV (9 kb) Human
 chr6
45,691,388 3728 1/7906 2246 45,691,417
Human HPV (18kb) Human
 chr10
A
B
X
11,700,488 5664 470 1972 11,700,451
HPV (37 kb) Human HPV (55 kb) Human segments
 chr19
6901 55,307,401 55,310,208 582
HPV (17 kb) Human (8 kb) HPV (49 kb)
 chrX
A
A
B
B
B
A
A
2986 145,708,231 145,716,514 3153
A
X
B
X
B
B

### Slide 4
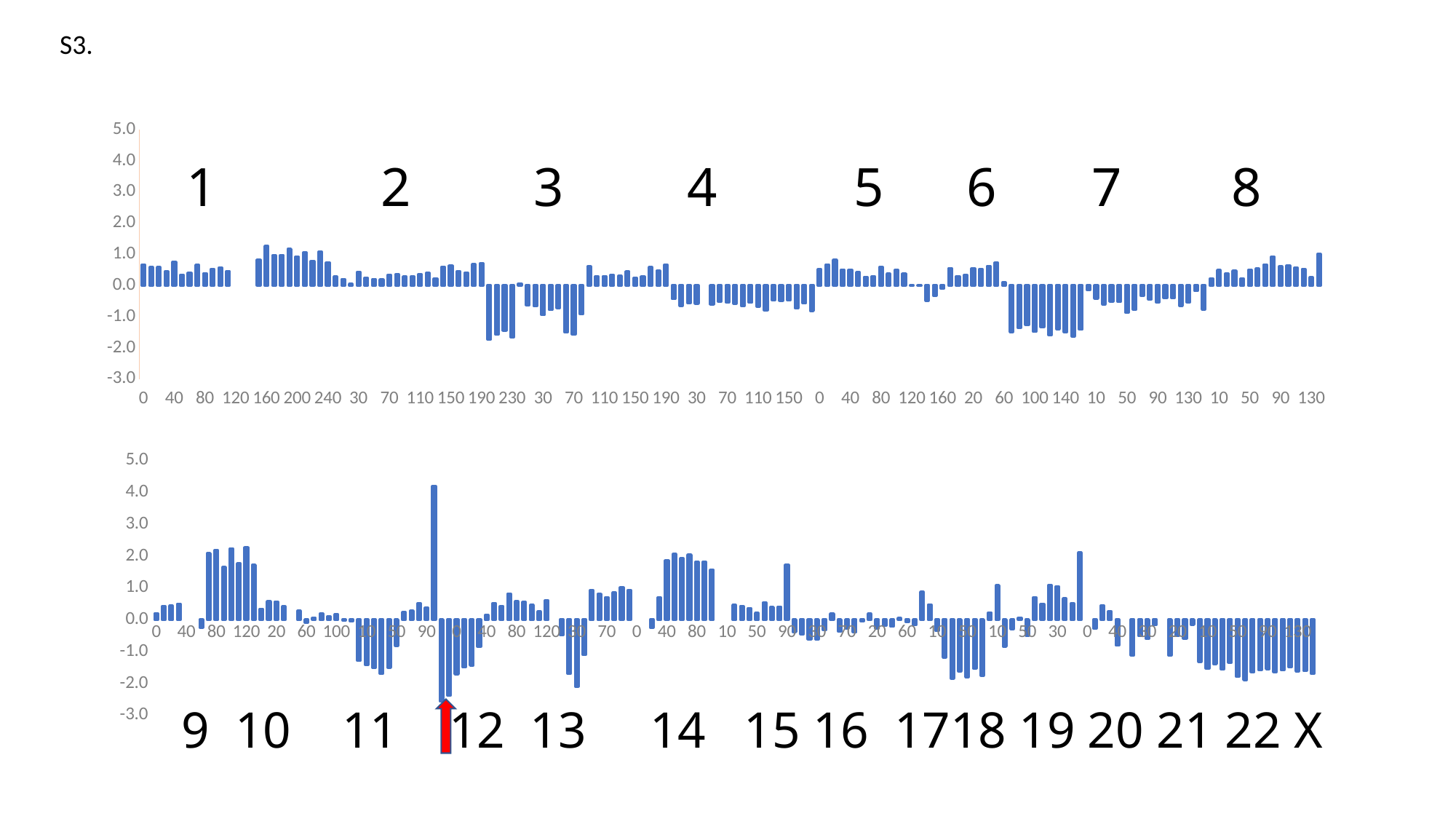

S3.
###### Chart
| Category | Z score |
|---|---|
| 9.9999999999999995E-7 | 0.660067315644761 |
| 10.000000999999999 | 0.574738835345594 |
| 20.000001000000001 | 0.574738835345594 |
| 30.000001000000001 | 0.4371122542179072 |
| 40.000000999999997 | 0.7481483275664806 |
| 50.000000999999997 | 0.32976352093831 |
| 60.000000999999997 | 0.3985768115021538 |
| 70.000000999999997 | 0.6518097207770993 |
| 80.000000999999997 | 0.3793090901442778 |
| 90.000000999999997 | 0.5169356712719654 |
| 100.000001 | 0.560976177232826 |
| 110.000001 | 0.4481223807081224 |
| 120.000001 | None |
| 130.000001 | None |
| 140.000001 | None |
| 150.000001 | 0.8142090865077708 |
| 160.000001 | 1.2601192093614781 |
| 170.000001 | 0.9573407308805656 |
| 180.000001 | 0.9655983257482271 |
| 190.000001 | 1.1665331341946505 |
| 200.000001 | 0.9133002249197057 |
| 210.000001 | 1.045421742802286 |
| 220.000001 | 0.7811787070371261 |
| 230.000001 | 1.0674419957827157 |
| 240.000001 | 0.7282689325147041 |
| 9.9999999999999995E-7 | 0.2719603568646814 |
| 10.000000999999999 | 0.1948894714331768 |
| 20.000001000000001 | 0.035242637325058945 |
| 30.000001000000001 | 0.42334959610513767 |
| 40.000000999999997 | 0.2361774457714823 |
| 50.000000999999997 | 0.19764200305573043 |
| 60.000000999999997 | 0.1783742816978537 |
| 70.000000999999997 | 0.33251605256086436 |
| 80.000000999999997 | 0.3517837739187411 |
| 90.000000999999997 | 0.27746542010978936 |
| 100.000001 | 0.27746542010978936 |
| 110.000001 | 0.3490312422961867 |
| 120.000001 | 0.390319216634493 |
| 130.000001 | 0.21966225603616074 |
| 140.000001 | 0.5857489618358092 |
| 150.000001 | 0.6242844045515619 |
| 160.000001 | 0.4481223807081224 |
| 170.000001 | 0.3985768115021538 |
| 180.000001 | 0.6683249105124217 |
| 190.000001 | 0.7013552899830664 |
| 200.000001 | -1.7263776011093415 |
| 210.000001 | -1.5667307670012243 |
| 220.000001 | -1.437361780741198 |
| 230.000001 | -1.6520592473003903 |
| 9.9999999999999995E-7 | 0.051757827060381296 |
| 10.000000999999999 | -0.6198598888427334 |
| 20.000001000000001 | -0.6556427999359324 |
| 30.000001000000001 | -0.9391535570589691 |
| 40.000000999999997 | -0.7767541913282973 |
| 50.000000999999997 | -0.7134459640095611 |
| 60.000000999999997 | -1.4951649448148265 |
| 70.000000999999997 | -1.5639782353786702 |
| 80.000000999999997 | -0.9143807724559848 |
| 90.000000999999997 | 0.6022641515711323 |
| 100.000001 | 0.28297048335489655 |
| 110.000001 | 0.2912280782225581 |
| 120.000001 | 0.32976352093831 |
| 130.000001 | 0.2939806098451117 |
| 140.000001 | 0.4371122542179072 |
| 150.000001 | 0.2334249141489295 |
| 160.000001 | 0.27746542010978936 |
| 170.000001 | 0.5719863037230404 |
| 180.000001 | 0.4618850388208904 |
| 190.000001 | 0.6552503853052916 |
| 9.9999999999999995E-7 | -0.4271826752639712 |
| 10.000000999999999 | -0.6418801418231637 |
| 20.000001000000001 | -0.55104659827889 |
| 30.000001000000001 | -0.5703143196367663 |
| 0 | None |
| 50.000000999999997 | -0.5978396358623038 |
| 60.000000999999997 | -0.5097586239405837 |
| 70.000000999999997 | -0.5372839401661212 |
| 80.000000999999997 | -0.5703143196367663 |
| 90.000000999999997 | -0.6501377366908249 |
| 100.000001 | -0.5400364717886752 |
| 110.000001 | -0.66114786318104 |
| 120.000001 | -0.79326938106362 |
| 130.000001 | -0.46296558635716983 |
| 140.000001 | -0.4822333077150462 |
| 150.000001 | -0.46571811797972346 |
| 160.000001 | -0.7217035588772226 |
| 170.000001 | -0.5648092563916587 |
| 180.000001 | -0.8097845707989424 |
| 9.9999999999999995E-7 | 0.505925544781751 |
| 10.000000999999999 | 0.6655723788898681 |
| 20.000001000000001 | 0.8087040232626637 |
| 30.000001000000001 | 0.5004204815366431 |
| 40.000000999999997 | 0.49491541829153585 |
| 50.000000999999997 | 0.4178445328600305 |
| 60.000000999999997 | 0.2637027619970206 |
| 70.000000999999997 | 0.28297048335489655 |
| 80.000000999999997 | 0.5774913669681476 |
| 90.000000999999997 | 0.3627939004089555 |
| 100.000001 | 0.4866578234238743 |
| 110.000001 | 0.36829896365406267 |
| 120.000001 | 0.0049647894769678375 |
| 130.000001 | -0.003292805390693729 |
| 140.000001 | -0.4767282444699386 |
| 150.000001 | -0.31708141036182114 |
| 160.000001 | -0.09412634893496744 |
| 170.000001 | 0.5472135191200564 |
| 9.9999999999999995E-7 | 0.2884755466000045 |
| 10.000000999999999 | 0.3215059260706492 |
| 20.000001000000001 | 0.5472135191200564 |
| 30.000001000000001 | 0.5086780764043046 |
| 40.000000999999997 | 0.6187793413064546 |
| 50.000000999999997 | 0.7178704797183896 |
| 60.000000999999997 | 0.08754073815358038 |
| 70.000000999999997 | -1.4786497550795037 |
| 80.000000999999997 | -1.3520333004420317 |
| 90.000000999999997 | -1.2611997568977578 |
| 100.000001 | -1.4758972234569505 |
| 110.000001 | -1.3355181107067091 |
| 120.000001 | -1.577740893491439 |
| 130.000001 | -1.3850636799126765 |
| 140.000001 | -1.4841548183246114 |
| 150.000001 | -1.6190288678297453 |
| 160.000001 | -1.4043314012705528 |
| 9.9999999999999995E-7 | -0.13816685489582692 |
| 10.000000999999999 | -0.40791495390609483 |
| 20.000001000000001 | -0.5950871042397499 |
| 30.000001000000001 | -0.5070060923180301 |
| 40.000000999999997 | -0.5180162188082449 |
| 50.000000999999997 | -0.8593301400049101 |
| 60.000000999999997 | -0.7602390015929749 |
| 70.000000999999997 | -0.31708141036182114 |
| 80.000000999999997 | -0.44369786499929353 |
| 90.000000999999997 | -0.5317788769210137 |
| 100.000001 | -0.40240989066098765 |
| 110.000001 | -0.39690482741588007 |
| 120.000001 | -0.63912761020061 |
| 130.000001 | -0.5290263452984596 |
| 140.000001 | -0.168444702743918 |
| 150.000001 | -0.7547339383478677 |
| 9.9999999999999995E-7 | 0.21415719279105278 |
| 10.000000999999999 | 0.4784002285562135 |
| 20.000001000000001 | 0.36829896365406267 |
| 30.000001000000001 | 0.4563799755757832 |
| 40.000000999999997 | 0.21140466116849918 |
| 50.000000999999997 | 0.4921628866689815 |
| 60.000000999999997 | 0.5279457977621805 |
| 70.000000999999997 | 0.6655723788898681 |
| 80.000000999999997 | 0.9133002249197057 |
| 90.000000999999997 | 0.5967590883260243 |
| 100.000001 | 0.6407995942868842 |
| 110.000001 | 0.560976177232826 |
| 120.000001 | 0.519688202894519 |
| 130.000001 | 0.24718757226169824 |
| 140.000001 | 1.0 |# 1 2 3 4 5 6 7 8
###### Chart
| Category | |
|---|---|
| 9.9999999999999995E-7 | 0.1948894714331768 |
| 10.000000999999999 | 0.4178445328600305 |
| 20.000001000000001 | 0.4288546593502457 |
| 30.000001000000001 | 0.49491541829153585 |
| 40.000000999999997 | None |
| 50.000000999999997 | None |
| 60.000000999999997 | -0.24001052493031616 |
| 70.000000999999997 | 2.08312616450505 |
| 80.000000999999997 | 2.1794647712944313 |
| 90.000000999999997 | 1.6399685732738962 |
| 100.000001 | 2.2345154037455064 |
| 110.000001 | 1.763832496288815 |
| 120.000001 | 2.2647932515935976 |
| 130.000001 | 1.7170394587054008 |
| 9.9999999999999995E-7 | 0.33251605256086436 |
| 10.000000999999999 | 0.5664812404779332 |
| 20.000001000000001 | 0.560976177232826 |
| 30.000001000000001 | 0.40408187474726176 |
| 40.000000999999997 | None |
| 50.000000999999997 | 0.2692078252421278 |
| 60.000000999999997 | -0.0803636908221983 |
| 70.000000999999997 | 0.05726289030548926 |
| 80.000000999999997 | 0.18112681332040806 |
| 90.000000999999997 | 0.09029326977613397 |
| 100.000001 | 0.16185909196253212 |
| 110.000001 | -0.014302931880908108 |
| 120.000001 | -0.039075716483892804 |
| 9.9999999999999995E-7 | -1.2832200098781876 |
| 10.000000999999999 | -1.4098364645156605 |
| 20.000001000000001 | -1.500670008059934 |
| 30.000001000000001 | -1.6850896267710356 |
| 40.000000999999997 | -1.4951649448148265 |
| 50.000000999999997 | -0.8042795075538348 |
| 60.000000999999997 | 0.22791985090382152 |
| 70.000000999999997 | 0.2692078252421278 |
| 80.000000999999997 | 0.5031730131591966 |
| 90.000000999999997 | 0.3627939004089555 |
| 100.000001 | 4.197070450626331 |
| 110.000001 | -2.5603946827431283 |
| 120.000001 | -2.3759750640320267 |
| 9.9999999999999995E-7 | -1.6988522848838044 |
| 10.000000999999999 | -1.4896598815697193 |
| 20.000001000000001 | -1.43185671749609 |
| 30.000001000000001 | -0.8455674818921414 |
| 40.000000999999997 | 0.13708630735954822 |
| 50.000000999999997 | 0.5141831396494118 |
| 60.000000999999997 | 0.4178445328600305 |
| 70.000000999999997 | 0.8142090865077708 |
| 80.000000999999997 | 0.5664812404779332 |
| 90.000000999999997 | 0.5472135191200564 |
| 100.000001 | 0.4536274439532288 |
| 110.000001 | 0.24718757226169824 |
| 120.000001 | 0.5995116199485787 |
| 9.9999999999999995E-7 | None |
| 10.000000999999999 | -0.47122318122483103 |
| 20.000001000000001 | -1.6878421583935892 |
| 30.000001000000001 | -2.0924643069089908 |
| 40.000000999999997 | -1.0932953279219786 |
| 50.000000999999997 | 0.9160527565422593 |
| 60.000000999999997 | 0.7976938967724477 |
| 70.000000999999997 | 0.6930976951154056 |
| 80.000000999999997 | 0.8417344027333084 |
| 90.000000999999997 | 1.0123913633316406 |
| 100.000001 | 0.9215578197873665 |
| 9.9999999999999995E-7 | None |
| 10.000000999999999 | None |
| 20.000001000000001 | -0.23725799330776218 |
| 30.000001000000001 | 0.6848401002477449 |
| 40.000000999999997 | 1.8574185714556426 |
| 50.000000999999997 | 2.052848316656959 |
| 60.000000999999997 | 1.9152217355292713 |
| 70.000000999999997 | 2.0308280636765286 |
| 80.000000999999997 | 1.8051204706271204 |
| 90.000000999999997 | 1.813378065494782 |
| 100.000001 | 1.5601451562198372 |
| 9.9999999999999995E-7 | None |
| 10.000000999999999 | None |
| 20.000001000000001 | 0.4536274439532288 |
| 30.000001000000001 | 0.4205970644825841 |
| 40.000000999999997 | 0.3572888371638475 |
| 50.000000999999997 | 0.2003945346782848 |
| 60.000000999999997 | 0.5334508610072886 |
| 70.000000999999997 | 0.390319216634493 |
| 80.000000999999997 | 0.40132934312470814 |
| 90.000000999999997 | 1.719791990327955 |
| 9.9999999999999995E-7 | -0.38038963768055734 |
| 10.000000999999999 | -0.44369786499929353 |
| 20.000001000000001 | -0.6088497623525186 |
| 30.000001000000001 | -0.603344699107411 |
| 40.000000999999997 | -0.3198339419843747 |
| 50.000000999999997 | 0.1756217500753001 |
| 60.000000999999997 | -0.358369384700127 |
| 70.000000999999997 | -0.2565257146656385 |
| 80.000000999999997 | -0.3886472325482185 |
| 9.9999999999999995E-7 | -0.0418282481064464 |
| 10.000000999999999 | 0.1756217500753001 |
| 20.000001000000001 | -0.2702883727784073 |
| 30.000001000000001 | -0.17670229761157957 |
| 40.000000999999997 | -0.20698014545967108 |
| 50.000000999999997 | 0.03799516894761254 |
| 60.000000999999997 | -0.06660103270942992 |
| 70.000000999999997 | -0.15743457625370363 |
| 80.000000999999997 | 0.8628371451728875 |
| 9.9999999999999995E-7 | 0.4701426336885519 |
| 10.000000999999999 | -0.34460672658735864 |
| 20.000001000000001 | -1.1703662133534836 |
| 30.000001000000001 | -1.836478866011492 |
| 40.000000999999997 | -1.6080187413395302 |
| 50.000000999999997 | -1.7896858284280779 |
| 60.000000999999997 | -1.5254427926629177 |
| 70.000000999999997 | -1.7456453224672177 |
| 9.9999999999999995E-7 | 0.21690972441360637 |
| 10.000000999999999 | 1.0674419957827157 |
| 20.000001000000001 | -0.8373098870244798 |
| 30.000001000000001 | -0.2785459676460688 |
| 40.000000999999997 | 0.05726289030548926 |
| 50.000000999999997 | -0.4849858393375998 |
| 9.9999999999999995E-7 | 0.6958502267379593 |
| 10.000000999999999 | 0.4921628866689815 |
| 20.000001000000001 | 1.0784521222729309 |
| 30.000001000000001 | 1.0371641479346245 |
| 40.000000999999997 | 0.6573147840222073 |
| 50.000000999999997 | 0.5031730131591966 |
| 60.000000999999997 | 2.1 |
| 9.9999999999999995E-7 | None |
| 10.000000999999999 | -0.26753584115585327 |
| 20.000001000000001 | 0.43986478584046085 |
| 30.000001000000001 | 0.25544516712935905 |
| 40.000000999999997 | -0.8015269759312815 |
| 9.9999999999999995E-7 | None |
| 10.000000999999999 | -1.1070579860347478 |
| 20.000001000000001 | -0.5015010290729225 |
| 30.000001000000001 | -0.5895820409946423 |
| 40.000000999999997 | -0.14917698138604207 |
| 9.9999999999999995E-7 | None |
| 10.000000999999999 | -1.1070579860347478 |
| 20.000001000000001 | -0.5015010290729225 |
| 30.000001000000001 | -0.5895820409946423 |
| 40.000000999999997 | -0.14917698138604207 |
| 9.9999999999999995E-7 | -1.3107453261037252 |
| 10.000000999999999 | -1.5337003875305792 |
| 20.000001000000001 | -1.3850636799126765 |
| 30.000001000000001 | -1.544710514020794 |
| 40.000000999999997 | -1.3437757055743702 |
| 50.000000999999997 | -1.7786757019378627 |
| 60.000000999999997 | -1.8887769668400132 |
| 70.000000999999997 | -1.6410491208101752 |
| 80.000000999999997 | -1.5749883618688851 |
| 90.000000999999997 | -1.544710514020794 |
| 100.000001 | -1.6355440575650677 |
| 110.000001 | -1.580493425113993 |
| 120.000001 | -1.4758972234569505 |
| 130.000001 | -1.6080187413395302 |
| 140.000001 | -1.599761146471869 |
| 150.000001 | -1.6924297110978455 | 9 10 11 12 13 14 15 16 1718 19 20 21 22 X

### Slide 5
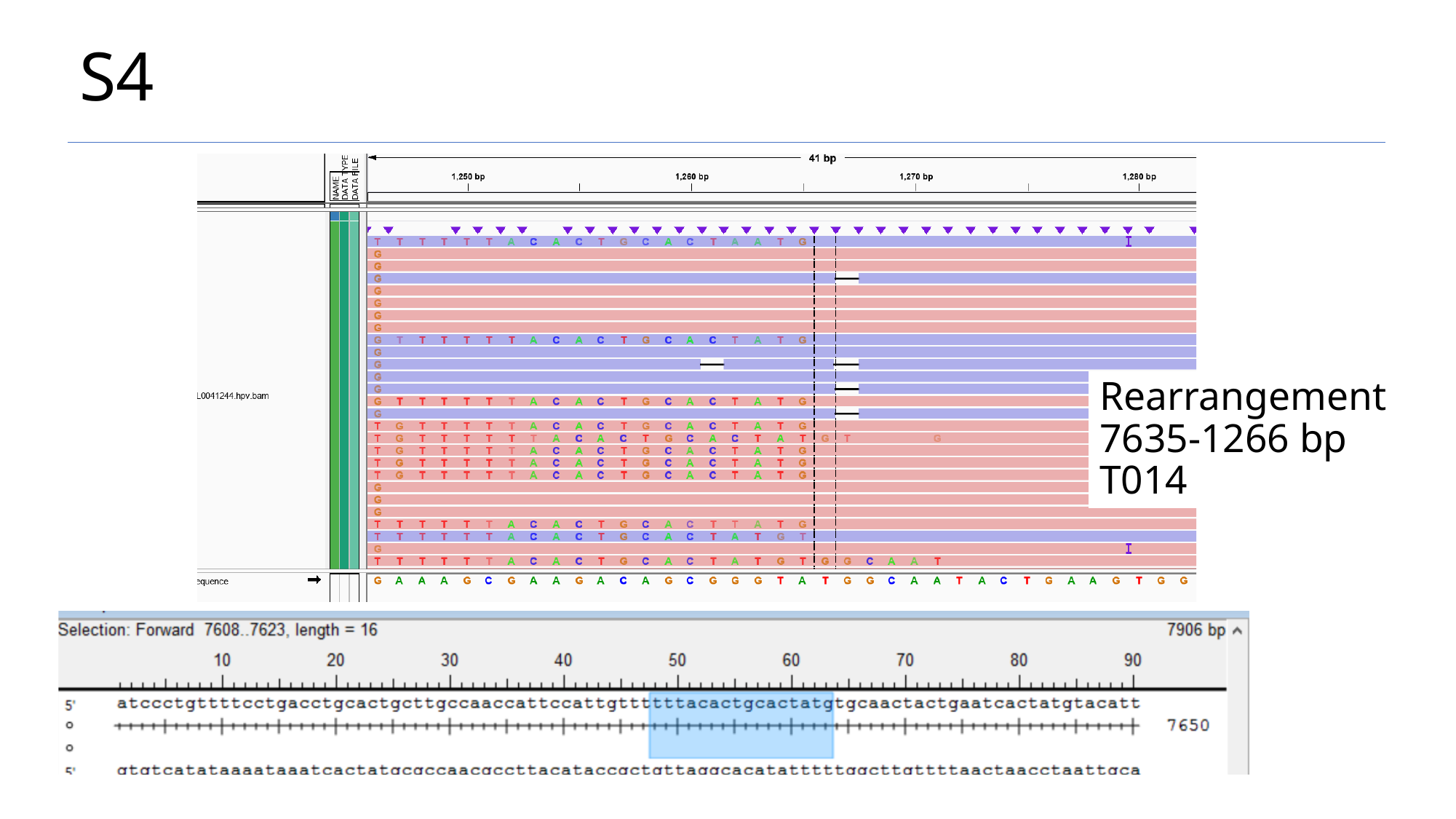

# S4
Rearrangement 7635-1266 bp
T014

### Slide 6
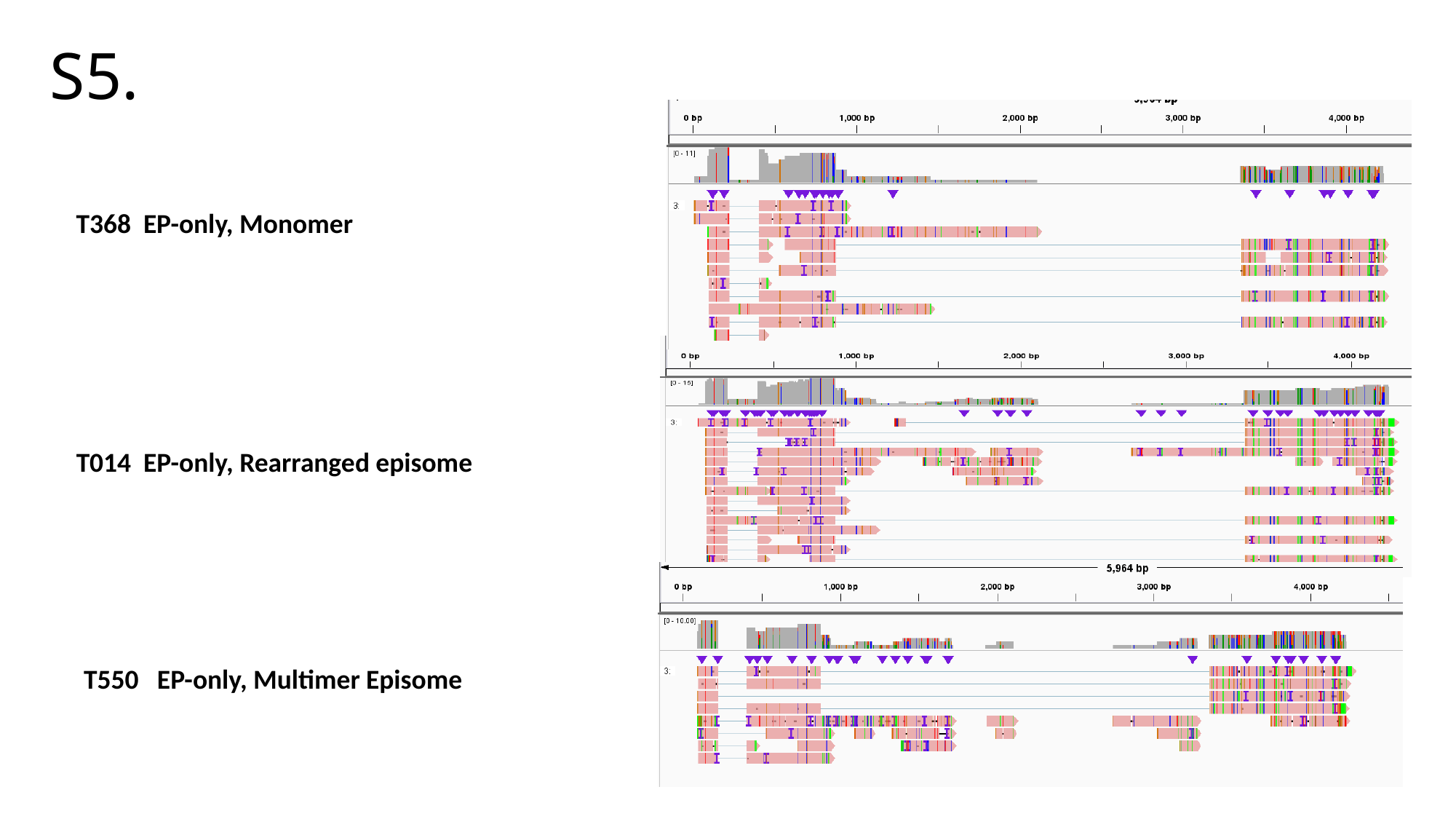

# S5.
T368 EP-only, Monomer
T014 EP-only, Rearranged episome
T550 EP-only, Multimer Episome
